# Phylogenetic reconciliation reveals extensive ancestral recombination in Sarbecoviruses and the SARS-CoV-2 lineage

**DOI:** 10.1101/2021.08.12.456131

**Authors:** Sumaira Zaman, Samuel Sledzieski, Bonnie Berger, Yi-Chieh Wu, Mukul S. Bansal

## Abstract

An accurate understanding of the evolutionary history of rapidly-evolving viruses like SARS-CoV-2, responsible for the COVID-19 pandemic, is crucial to tracking and preventing the spread of emerging pathogens. However, viruses undergo frequent recombination, which makes it difficult to trace their evolutionary history using traditional phylogenetic methods. Here, we present a phylogenetic workflow, virDTL, for analyzing viral evolution in the presence of recombination. Our approach leverages reconciliation methods developed for inferring horizontal gene transfer in prokaryotes, and, compared to existing tools, is uniquely able to identify ancestral recombinations while accounting for several sources of inference uncertainty, including in the construction of a strain tree, estimation and rooting of gene family trees, and reconciliation itself. We apply this workflow to the *Sarbecovirus* subgenus and demonstrate how a principled analysis of predicted recombination gives insight into the evolution of SARS-CoV-2. In addition to providing confirming evidence for the horseshoe bat as its zoonotic origin, we identify several ancestral recombination events that merit further study.

## Introduction

Phylogenetic analysis of the first available sequence from Wuhan, China placed SARS-CoV-2 in the *Sarbecovirus* subgenus of *Betacoronavirus* [49], and several subsequent studies have investigated its evolutionary origins [1,7]. SARS-CoV-2 shares 96% sequence similarity to bat *Sarbecovirus* RaTG13, and the two viruses form a clade distinct from other SARS-related coronaviruses, suggesting that the SARS-CoV-2 lineage may have its zoonotic origins in bats [51]. Pangolins have also been suggested as possible hosts [25], though later studies have shown that, while pangolins are natural reservoirs of *Betacoronaviruses*, SARS-CoV-2 likely did not evolve directly from pangolin coronavirus [27]. Language models have also shown that SARS-CoV-2 is “semantically” closest to bat and next to pangolin [19]. Such analyses are of biological and societal interest, as identifying the source of the virus may help inform future outbreaks of viruses with zoonotic origins.

Many viruses, including coronaviruses, undergo frequent recombination [13, 31], which complicates phylogenetic analysis [33]. Moreover, phylogenetic inference is susceptible to several sources of uncertainty, many of which are exacerbated by recombination between viral genomes. Thus, a common step in the study of viral evolution is to infer recombination, which is commonly done by enumerating triplets of strains and analyzing their sequence similarity [28, 30]. This approach works well when recombination occurs infrequently (relative to the rate of evolution) and mostly between extant strains (Figure 1a). However, as the number of strains grows and recombination occurs multiple times within a lineage, recombination becomes difficult to infer from sequence similarity alone (Figure 1b).

**Fig. 1:**
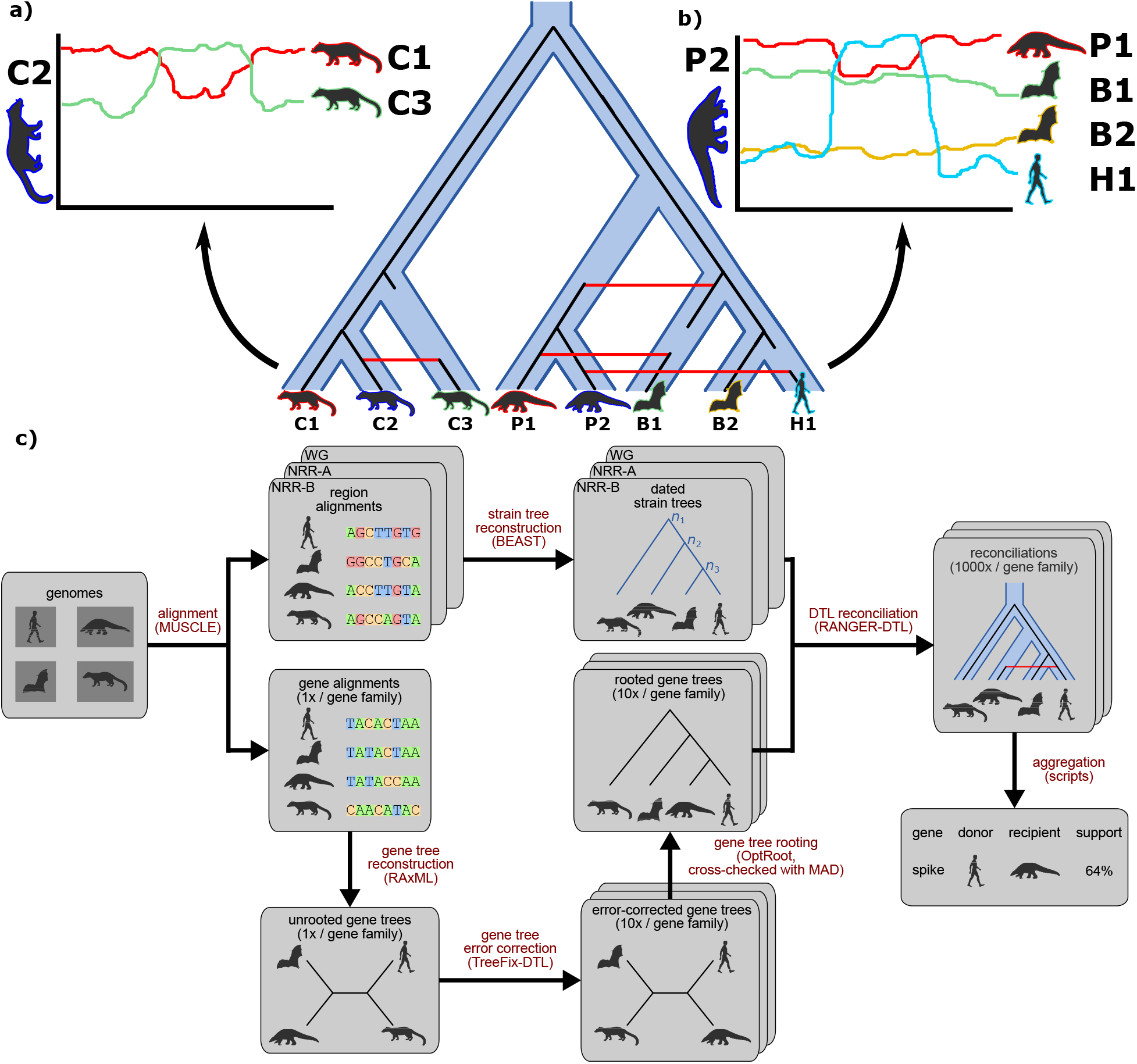
virDTL enables inference of ancestral recombination. **(a)** Commonly-used tools such as Simplot and RDP are well-suited to inferring recent recombinations between strains of interest, where the recombination signal is clear in the sequence similarity profile. **( b)** However, in cases where recombination has occurred between ancestral strains, and multiple recombinations have occurred in a single lineage, if becomes significantly more difficult to disentangle the sequence similarity signal to infer all recombinations. **(c)** Our model-based computational protocol, virDTL, takes into account the entire evolutionary history of a gene family, including several sources of inference uncertainty. A credible strain tree is estimated using non-recombinant regions of the genome, and multiple gene tree candidates are inferred and error-corrected and reconciled against the strain tree to infer HGTs. In addition to accounting for gene tree topological and rooting uncertainty, we reconcile the same gene tree and species tree multiple times to capture the full landscape of uncertainty in inferring recombination.

Recombination in viruses is similar to gene conversion in that it generally results in the one-sided transfer of genetic material from a donor genome to a recipient genome, rather than an “exchange” of genetic material between the two recombining genomes [34]. Thus, we reasoned that methods for studying horizontal gene transfer (HGT) in prokaryotes could potentially be useful for inferring recombination in viruses. Despite advances in horizontal gene transfer (HGT) detection (*Discussion*), these methods have rarely been used to study viral genome evolution or recombination.

In this work, we describe a step-by-step computational protocol, virDTL, for analyzing viral evolution in the presence of recombination. virDTL newly leverages Duplication-Transfer-Loss (DTL) reconciliation, a powerful computational technique used to study horizontal gene transfer in prokaryotes [2–4, 8, 9, 11, 15, 20, 24, 26, 37–39, 41, 43–46], to gain insights into viral evolution and recombination (Figure 1c). In addition, virDTL addresses common sources of HGT inference error and uncertainty under recombination by carefully constructing the strain tree and by using resampling and error-correction methods. virDTL addresses some of the key difficulties traditionally associated with viral evolutionary analysis, such as systematic, large-scale identification of ancestral recombination events and precise phylogenetic identification of the recombining strains, and can help virologists and epidemiologists better understand viral evolution and easily infer recombination events.

We demonstrate the utility of virDTL by using it to investigate viral recombination in the *Sarbecovirus* subgenus. Specifically, we ran virDTL on 54 *Sarbecovirus* genomes from 4 host species, including the novel coronavirus SARS-CoV-2, and assessed its ability to recover recombinations between leaf strains and discover new ancestral recombinations (*Results*). We identify 226 plausible leaf-to-leaf (i.e., between sampled strains) and 362 plausible ancestral HGTs across all gene families, and identify 8 well-supported HGTs of potential relevance to SARS-CoV-2 evolution, including 3 in the well-studied Spike and Nucleocapsid proteins. We use the popular sequence similarity tool SimPlot [28] to validate our protocol on a subset of leaf-to-leaf HGTs and explore several case studies where our DTL-reconciliation-based approach enables inference of viral recombination. Among other results, our analysis supports the previously-proposed hypothesis that similarity between the SARS-CoV-2 and pangolin strains arose due to a recombination between the ancestor of SARS-CoV-2 and RaTG13 and an ancestral pangolin viral strain [7].

Lastly, we identify and discuss the strengths and limitations of the proposed reconciliation-based approach, contrast virDTL with widely-used sequence-similarity based approaches such as SimPlot [28] and RDP [30], and compare our protocol against two recent approaches used to investigate recombination in coronaviruses using phylogenetic reconciliation, developed in parallel and independently from this work [14, 29] (*Discussion*).

## Results

### Overview of virDTL

The virDTL protocol is designed to infer recombination in viruses while minimizing the impact of key sources of error (Figure 1c). We briefly describe the key steps below.

#### 1. Strain tree reconstruction and selection

Since viruses are frequently impacted by substantial recombination, virDTL first identifies non-recombinant or minimally-recombinant genomic regions that could be used to reconstruct credible strain trees. It then further analyzes candidate strain trees to identify a single, minimally recombinant strain tree. virDTL uses BEAST [42] for candidate strain tree reconstruction.

#### 2. Gene tree reconstruction and error-correction

Gene trees are often impacted by phylogenetic reconstruction error and uncertainty due to lack of sufficient phylogenetic signal. virDTL minimizes the downstream impact of such error and uncertainty by error-correcting the gene tree topologies to match the strain tree unless the sequence data confidently supports incongruence. virDTL uses RAxML [40] for initial gene tree construction and TreeFix-DTL [5] for error-correction. virDTL further accounts for topological uncertainty by sampling multiple error-corrected gene trees per gene family for reconciliation analysis.

#### 3. Gene tree rooting

Gene trees reconstructed using standard phylogenetic approaches are unrooted and must be rooted prior to reconciliation analysis. Since there is often uncertainty in rooting gene trees, virDTL uses multiple gene tree rooting approaches and assess how the resulting differently rooted gene trees affect support for final evolutionary inferences. virDTL uses OptRoot [4] and Minimum Ancestor Deviation (MAD) rooting [47], with OptRoot as the primary rooting method.

#### 4. Phylogenetic reconciliation analysis

To account for ambiguity or uncertainty in phylogenetic reconciliation, virDTL randomly samples many optimal reconciliations per gene tree and aggregates inferences across both reconciliation samples and gene tree samples to identify only well-supported HGTs for each gene family. virDTL uses RANGER-DTL [4] to sample optimal reconciliations.

#### 5. Strain tree dating and evaluation of HGTs

virDTL dates the strain tree so that any HGTs inferred can be evaluated for time-consistency among the participating strains, and performs additional analysis to determine if the detected HGTs support the inference of larger recombination events. virDTL uses BEAST [42] to perform strain tree dating.

Full details of the virDTL protocol as well as its specific application to our *Sarbecovirus* dataset appear in *Methods*. Scripts implementing several aspects of the virDTL protocol, along with step-by-step instructions, are freely available at https://github.com/suz11001/virDTL.

### Recombination occurs frequently in Sarbecoviruses

The *Sarbecovirus* genome comprises 4 well-characterized structural proteins which construct the viral Spike (S), Envelope (E), Membrane (M), and Nucleocapsid (N), and 7 open reading frames which act as accessory factors (Figure 2b). The largest open reading frame, ORF1ab, comprises the replicase-transcriptase complex displayed as two polyproteins (ORF1a and ORF1b), which synthesize 16 non-structural proteins by three viral proteases [17,22,23]. The smaller open reading frames, near the 3’ end of the genomes, encode proteins hypothesized to interact with a diverse array of host biological pathways [16].

**Fig. 2:**
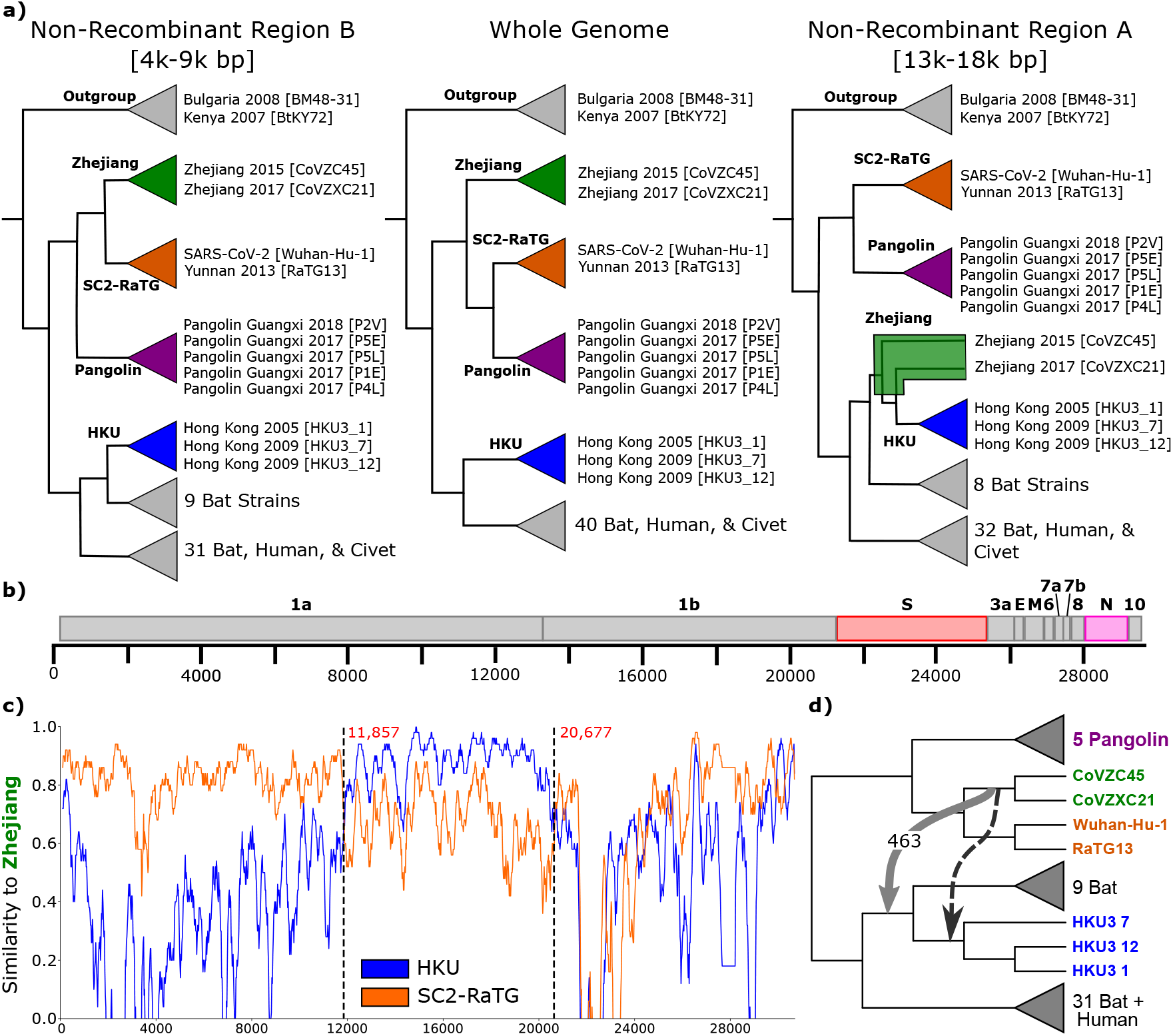
Overview of *Sarbecovirus* genome evolution. **(a)** We reconstructed three candidate strain trees from the whole genome and two putative non-recombinant regions A (13,000 - 18,000 base pairs) and B (4,000 - 9,000 base pairs). Their topologies differ substantially, especially in the SARS-CoV-2 lineage, which suggests that the evolution of SARS-CoV-2 was impacted by recombination. We define 4 clades, *Zhejiang* (green), *SC2-RaTG* (orange), *Pangolin* (purple) and *HKU* (blue) and show the tree inferred using each region of the genome. **(b)** The *Sarbecovirus* genome comprises 4 well-characterized structural proteins which construct the viral Spike (S), Envelope (E), Membrane (M), and Nucleocapsid (N), as well as several open reading frames which encode accessory factors. The Spike and Nucleocapsid genes are highlighted in red and pink, respectively, as they appear in several ancestral recombinations (Figure 5). **(c)** Sequence similarity along the genome using SimPlot. Using *Zhejiang* clade sequences as query, we compare with the *SC2-RaTG* and *HKU* clades. For the majority of the genome, *SC2-RaTG* is more similar to *Zhejiang*. Between 11,857 and 20,677 base pairs, *HKU* is more similar. **(d)** We find evidence of an HGT from the ancestor of the *Zhejiang* clade to an ancestor of the *HKU* clade in ORF-1ab. This recombination (light gray) explains the signal shown in the NRR-A tree **(a)** and SimPlot **(c)** and is not consistent with the dating of the phylogeny (Figure S2). However, it is not uncommon for inferred HGTs to be off by a single branch due to gene tree estimation uncertainty. A time-consistent HGT to the ancestor of the three *HKU* strains (darker gray) similarly explains the signal.

We construct 3 candidate strain trees using the whole genome and two putative non-recombinant regions A and B (Figure 2a; *Methods*). Due to potential recombination in non-recombinant region A (Figure 2c,d; *Methods*), we select the non-recombinant region B tree against which to reconcile our gene trees.

We account for topological uncertainty and reconciliation uncertainty by reconstructing 10 error-corrected gene trees per gene family and, for each rooting, sampling 100 optimal DTL reconciliations for each gene tree. Thus, an inferred HGT can have a maximum support of 1000. In this analysis, we consider an HGT to be *supported* if it is found in at least 100 samples, which corresponds to HGTs whose support values are in the 95^th^ percentile. Note that support quantile thresholds were computed by aggregating results across undirected donor-recipient pairs, and so are more stringent than for single directed HGT events. Using OptRoot-rooted gene trees, we identified 588 supported HGT events across all gene families (Supplementary Table S3), which includes 78 with at least 500 support (99^th^ percentile), and 25 with at least 808 support (99.5^th^ percentile). Of the supported HGTs, 226 are leaf-to-leaf, 115 are ancestor-to-ancestor, and 247 involve an ancestral node and a leaf. As a different rooting of the gene tree may affect the inferred events, we verified that, of the 588 HGTs, 441 are also supported using MAD rooting. Of the 78 HGTs with support in the 99^th^ percentile, 71 were also supported using MAD rooting, including all 25 HGTs with support in the 99.5^th^ percentile.

We verified that most HGTs in the 99^th^ percentile are consistent with temporal constraints implied by the divergence times estimated on our strain tree. Note that HGTs that go forward in time can be temporally consistent due to the existence of unsampled strains [10], but HGTs cannot go backward in time. Specifically, we found that of the 78 HGTs with support in the 99^th^ percentile, 66 are consistent with the dating implied by the strain tree, including 24 out of 25 HGTs with support in the 99.5^th^ percentile. Seven additional HGTs would be time-consistent if the donor or recipient were shifted by one branch, leaving only 5 of the 78 events as fully inconsistent. We note that both estimating divergence time and identifying donors and recipients of HGT events can be error-prone, and some inconsistency is therefore expected.

While a detailed analysis of all putative HGT events is beyond the scope of this work, we highlight 8 HGTs involving the SARS-CoV-2 lineage, including a recombination in the Spike protein between an ancestor of Pangolin viral strains and an ancestor of SARS-CoV-2 and Bat CoV RaTG13. We also validate a subset of inferred leaf-to-leaf HGTs using the sequence-based recombination detection tool SimPlot [28] and highlight additional case studies for ancestral HGTs and HGTs with ambiguous direction. In addition, we assess the feasibility of using inferred HGTs to detect larger recombination events spanning multiple genes.

Time-consistent HGTs with ancestral recipients and greater than 500 support are shown in Figure S2. We provide a full list of all inferred HGTs (Supplementary Table S2) as well as the full strain tree with all internal nodes labeled (Supplementary Figure S1).

### Recombination affects the SARS-CoV-2 lineage

Given the interest in understanding SARS-CoV-2 evolution, we used virDTL to search for recombinations involving the SARS-CoV-2 lineage. We inferred six highly-supported HGTs using the default OptRoot-rooted gene trees, and two additional highly-supported HGTs using MAD-rooted gene trees (Figure 3a). Among these events, at least two are transfers into the SARS-CoV-2 lineage and at least one is a transfer from the SARS-CoV-2 lineage; a clear direction of transfer could not be inferred for the remaining five HGTs. All HGTs involved ancestors of SARS-CoV-2 rather than SARS-CoV-2 itself.

**Fig. 3:**
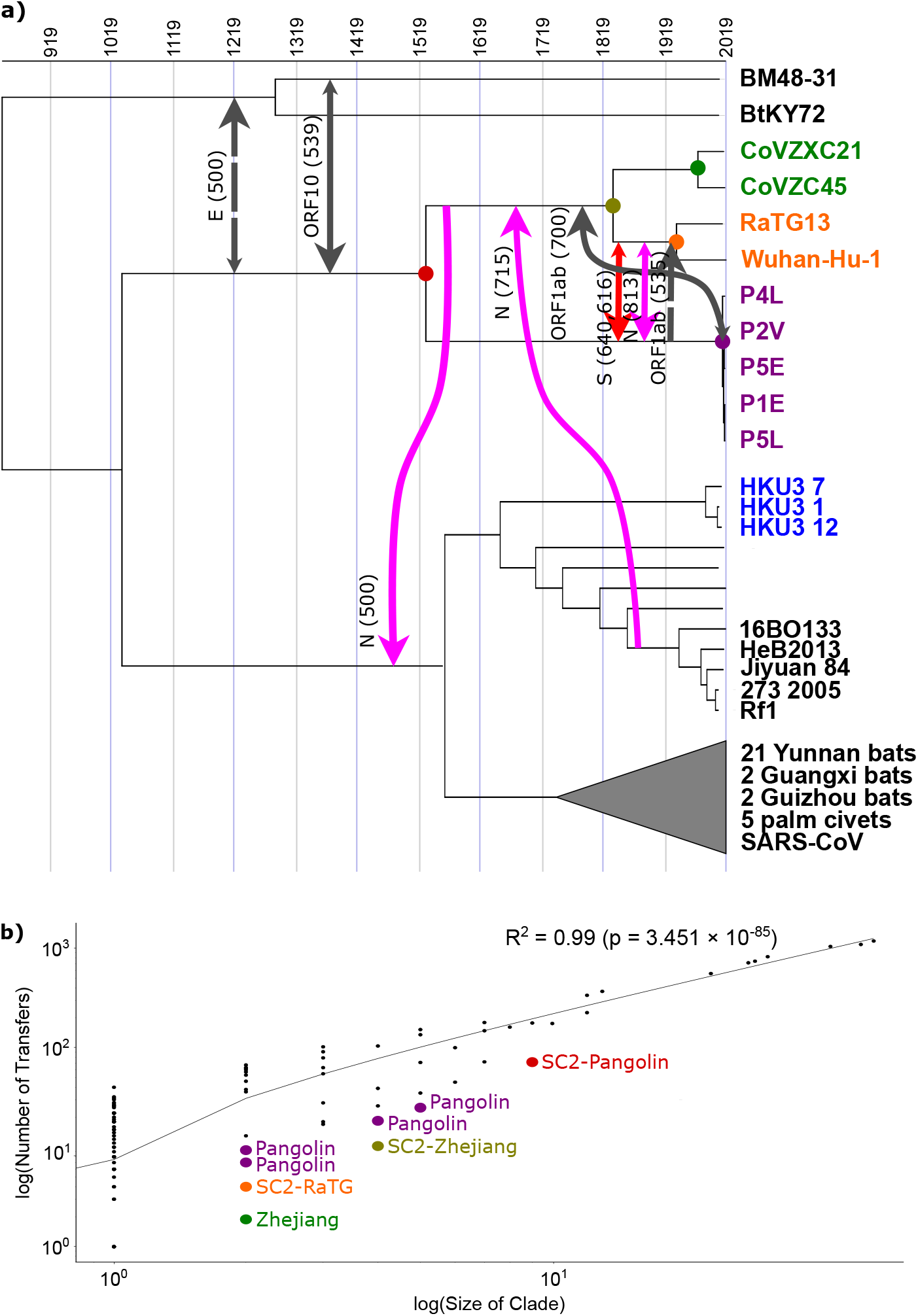
HGTs involving the SARS-CoV-2 lineage. **(a)** We inferred eight highly supported (with a support of at least 500) HGTs which involve an ancestor of SARS-CoV-2 [Wuhan-Hu-1]. Support values are shown for the OptRoot-rooted gene trees (solid lines) or MAD-rooted gene trees (dashed lines), with one transfer (Spike) inferred using both rootings. Smaller arrow heads indicate there exists an HGT with at least 100 support in the reverse direction using gene trees rooted with either method, suggesting directional uncertainty. **(b)** We found a strong correlation between the size of a clade and the number of HGTs in that clade (Pearson’s *R*^2^ = 0.99). However, for every ancestral strain in the SARS-CoV-2 lineage and related clades, the number of HGTs in that clade is much lower than would be expected for their size. This paucity of HGTs is likely due to sampling effects, as these strains are more distantly related to the rest of the *Sarbecovirus* strains in the analysis.

Most prominently, we found evidence for recent recombination in the Spike gene family, between the ancestor of Wuhan-Hu-1 (i.e., SARS-CoV-2) and RaTG13 and the ancestor of the pangolin strains. Ignoring directionality, this time-consistent transfer has a support of 1000 using both OptRoot- and MAD-rooted gene trees; that is, it is supported by every gene tree and reconciliation. However, support was roughly evenly split between the two directions, with OptRoot showing support of 640 (360) and MAD showing support of 616 (384) for a transfer from (to) the ancestor of Wuhan-Hu-1 and RaTG13. We discuss a possible cause of this directional uncertainty later in the manuscript (*Case Study: Spike and Nucleocapsid HGTs*). Despite directional uncertainty, this HGT supports the previously-proposed hypothesis that similarity between SARS-CoV-2 and pangolin strains arose due to recombination rather than pangolins being a possible host [7].

We also found evidence for three recombinations in the Nucleocapsid gene family. One of these HGTs is similar to the Spike gene HGT discussed above. When using OptRoot (MAD), this HGT has a support of 813 (456) from the ancestor of Wuhan-Hu-1 and RaTG13 to the ancestor of the pangolin strains and a support of <100 (444) in the reverse direction. Thus, we view the direction of this HGT as uncertain even though our default results using OptRoot suggest that it occurred from the SARS-CoV-2 lineage to the ancestor of the pangolin strains. The second Nucleocapsid HGT is a transfer from the ancestor of South Korean, Hebei, Henan, and Hubei bat strains to the ancestor of Wuhan-Hu-1 and Zhejiang strains. This HGT is time-consistent within one branch and is highly supported using OptRoot (715) but not MAD (< 100). The third Nucleocapsid HGT is a transfer from the ancestor of Wuhan-Hu-1 and Zhejiang strains to the ancestor of SARS-CoV, several Hong Kong and other Asian bat strains, and civet strains. While this HGT is time-consistent, it is again only supported by OptRoot (500) and not MAD (< 100).

We also found evidence for recombination in several other gene families. One of these is a time-consistent HGT affecting the ORF1ab gene and is similar to the previously observed HGTs in the Spike and Nucleocapsid genes, between between the ancestor of Wuhan-Hu-1 and RaTG13 and the ancestor of the pangolin strains. While it is not supported by OptRoot-rooted gene trees, it has an undirected support of 1000 using MAD-rooted gene trees. However, this HGT also shows directional uncertainty, with a roughly evenly split support of 535 and 465 in the two directions. Additionally, we found two time-consistent transfers between the outgroup of Bulgaria and Kenyan bat strains and the ancestor of Wuhan-Hu-1 and pangolin strains. One of these HGTs occurs in the ORF10 gene and also shows directional uncertainty, with OptRoot showing support of 539 (261) and MAD showing support of 378 (122) for a transfer to (from) the SARS-CoV-2 lineage. A similar HGT occurs in the Envelope gene, with OptRoot showing support of 147 (243) and MAD showing support of 500 (<100) for a transfer from (to) the SARS-CoV-2 lineage. Lastly, we found a time-inconsistent HGT in ORF1ab from the ancestor of the four pangolin strains to the ancestor of Wuhan-Hu-1 and the Zhejiang strains. We note that we also find several other transfers with lower but still substantial support (≥ 100) that might warrant further investigation (Supplementary Table S2).

Interestingly, by analyzing the donors and recipients of our full list of 588 HGTs supported using OptRoot-rooted gene trees, we found that the ancestors and nearby relatives of the SARS-CoV-2 [Wuhan-Hu-1] genome uniformly undergo recombination less often than the rest of the *Sarbecovirus* subgenus (Figure 3b). However, this observation may be an artifact of sampling effects caused by the relatively small number of strains in this clade, and because of the low overall diversity among these strains.

### Reconciliation recovers HGTs between leaf strains

After finding evidence for recombination involving the SARS-CoV-2 lineage, we expanded our analysis to the entire *Sarbecovirus* subgenus. Our analysis identified 226 supported leaf-to-leaf HGTs (≥ 100 support) and 35 highly supported leaf-to-leaf HGTs (≥ 500 support). Among the 35 highly supported HGTs, 34 were intra-host HGTs between strains from the same host type (e.g. bat to bat, pangolin to pangolin, etc.), and one was an inter-host HGTs between strains from different host types (human SARS-CoV to civet C010 in ORF7a). Such leaf-to-leaf HGTs can be orthogonally verified through a SimPlot analysis by choosing the recipient, donor, and sister strains of both recipient and donor, as demonstrated through the following case study.

#### Case Study: Spike gene HGT between strains from bats

We identified an HGT in the Spike gene between the strains Guangxi 2004 [Rp3, donor] and Hubei 2004 [Rm1, recipient] with a support of 1000 (Figure 4a). For the SimPlot analysis, we selected GX2013 as the sister of Rp3 and HuB2013 as the sister of Rm1. When querying the genome of Rp3, we see high similarity with its sister GX2013 throughout the entire length of its genome with the exception of the Spike gene region, where Rp3 is most similar to the recipient Rm1 (Figure 4b). Reciprocally, when querying the genome of Rm1, we see that sequence similarity with HuB2013 decreases in the region encompassing the Spike gene while sequence similarity with Rp3 increases (Figure 4c). These findings are consistent with a hypothesis in which Rm1 received Rp3’s copy of the Spike gene. Additionally, we observe that Rm1 continues to remain highly similar to Rp3 even beyond the boundary of the Spike gene. This observation could indicate a larger multi-gene recombination event, which was not detected in our reconciliation-based analysis.

**Fig. 4:**
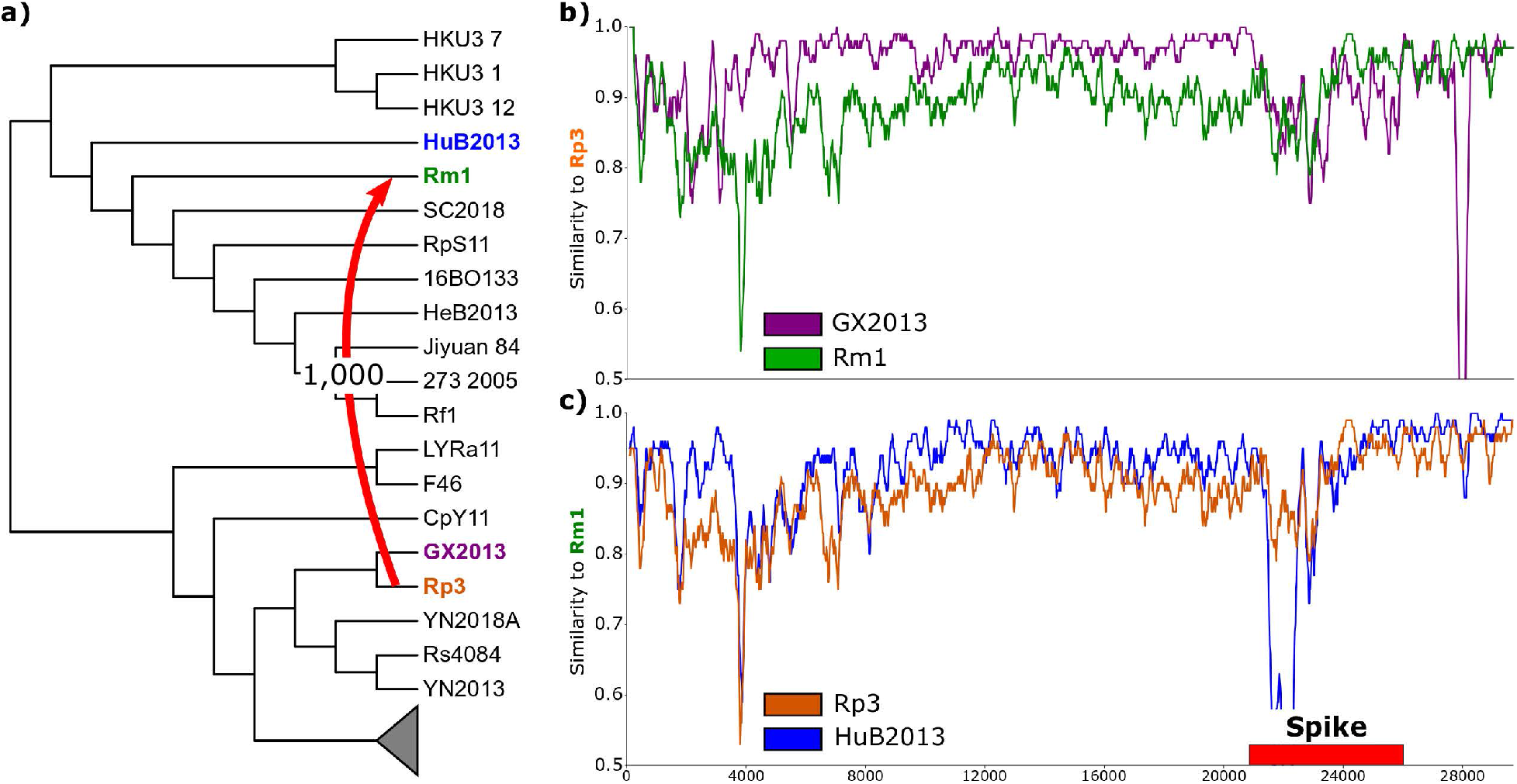
Highly supported leaf-to-leaf HGTs are consistent with sequence similarity. We present a case study of a leaf-to-leaf HGT between the donor Rp3 (orange) and recipient Rm1 (green). **(a)** The inferred HGT from Rp3 to Rm1 in the Spike gene has a support of 1000. **(b)** Sequence similarity of the donor Rp3 to its sibling GX2013 (purple) and the recipient Rm1. Rp3 and GX2013 are highly similar throughout the length of the genome, and Rm1 is more divergent throughout but equally similar in the Spike region. **(c)** Sequence similarity of the recipient Rm1 to its sibling HuB2013 (blue) and the donor Rp3. Rm1 and HuB2013 are highly similar throughout the length of the genome *except* in the Spike region, where Rm1 has received genetic material from Rp3. Thus, Rp3 and Rm1 are more similar in the Spike region.

To further assess the accuracy of recombination events inferred through virDTL, we performed similar SimPlot analyses using the donor, recipient, and recipient-sister strains to orthogonally verify each of the five other highly supported HGTs identified by virDTL in the Spike gene. The Spike gene is a good candidate for such a SimPlot analysis since it is sufficiently long for recombinations to be easily visible and interpretable. We find a clear signal for recombination in the Spike gene for each of the five cases (Figure S3). In three of the cases (F46 to Rf4092, Rs4081 to YN2018D, and Anlong-103 to YN2013), the recombination affects predominantly the Spike gene region. In the other two cases (Jiyuan 84 to HeB2013 and Rs9401 to Rs7327), the donor sequence is more similar to the recipient sequence along the majority of the genome. Possible explanations include that the Spike gene HGT may be part of a larger recombination event, that the Spike gene HGT may be an artifact of incorrect placement of the affected strains in the strain tree, or that the recipient sister sequence has undergone rapid evolution. Analysis of the sequences and dated species tree suggest the latter is more likely for the transfer from Jiyuan 84 to HeB2013, while the transfer from Rs9401 to Rs7327 is more likely due to a multi-gene recombination.

### Reconciliation reveals new ancestral HGTs

While we validate our approach on leaf-to-leaf transfers, the virDTL protocol also enables the inference of ancestral recombination. Our analysis identified 115 supported and 11 highly supported ancestor-to-ancestor HGTs, 113 supported and 14 highly supported ancestor-to-leaf HGTs, and 134 supported and 18 highly supported leaf-to-ancestor HGTs. While ancestor-to-ancestor and leaf-to-ancestor HGTs must correspond to an ancestral HGT, ancestor-to-leaf HGTs may in fact be an HGT from an unsampled leaf to a sampled leaf. Among the 11 highly supported ancestor-to-ancestor HGTs, 5 involve the SARS-CoV-2 lineage, 2 were intra-host HGTs between clades that contain the same host type, and 9 were inter-host HGTs between clades that contain different host types. Since SimPlot compares sequences of extant strains, it is more difficult to verify ancestral recombination events through the kind of external analysis demonstrated above for leaf-to-leaf HGTs. Despite this limitation, in the following case study, by using appropriately chosen descendants of the ancestral donor and recipient, we demonstrate that observed genomic sequence similarity is consistent with the inferred ancestral recombination. However, we note that *post facto* investigation of inferred ancestral HGTs using sequence similarity is more feasible than discovery of such HGTs from similarity alone.

#### Case Study: Spike and Nucleocapsid HGTs

As previously reported, we identified highly supported HGTs in the Spike and Nucleocapsid genes between the common ancestor of Wuhan-Hu-1 (i.e., SARS-CoV-2) and RaTG13 (hence *SC2-RaTG*) and the common ancestor of the pangolin strains (*Pangolin*). The Spike gene HGT shows a support value of 640 from *SC2-RaTG* to *Pangolin* and 360 in the reverse direction when using OptRoot for gene tree rooting, and 616 and 384, respectively, when using MAD rooting. Likewise, the Nucleocapsid HGT has a support of 813 from *SC2-RaTG* to *Pangolin* using OptRoot rooting but 444 in the reverse direction when using MAD rooting. Thus, while the analysis clearly shows that recombination occurred between *SC2-RaTG* and *Pangolin* in both the Spike and Nucleocapsid genes, the direction of these HGTs cannot be unambiguously inferred through our analysis. This ambiguity in direction inference is the result of a lack of resolution in the species tree, such that an HGT in either direction between *SC2-RaTG* and *Pangolin* may be able to explain the corresponding gene tree topologies. Nonetheless, in this case study we demonstrate how it may sometimes be possible to use sequence similarity to additionally support inferred ancestral HGTs. The SimPlot analysis below also suggests that both the Spike and Nucleosapsid genes may have been transferred in a single recombination event.

Using *SC2-RaTG* as the query, we found that the most closely related strain, the ancestor of CoVZC45 and CoVZXC21 (*Zhejiang*) is more similar for much of the genome (Figure 5a), but that *Pangolin* becomes more similar for both the Spike and regions of the Nucleocapsid gene (Figure 5c). This finding is consistent with prior literature indicating similarity between the pangolin strains and the SARS-CoV-2 [Wuhan-Hu-1] genome in the Spike protein [25, 50]. However, our analysis suggests that this similarity can be accounted for by a recombination between the ancestor of Wuhan-Hu-1 and RaTG13 and the ancestor of the pangolin strains, which is consistent with the findings of Boni et al. [7]. The elevated similarity in parts of the Nucleocapsid sequence is likely a result of the same recombination event.

**Fig. 5:**
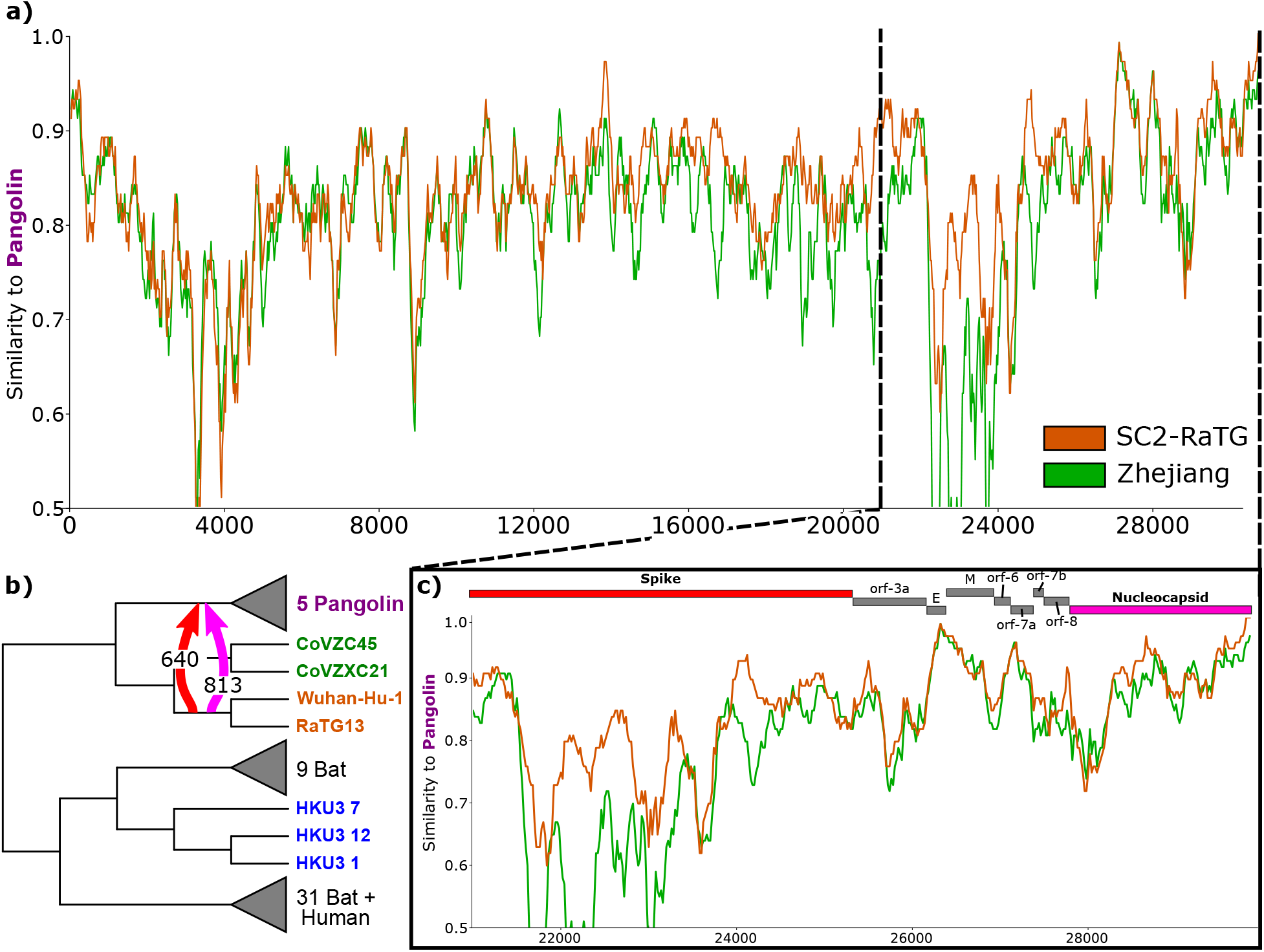
Ancestral HGTs are consistent with sequence similarity but difficult to discover from similarity alone. We present a case study of an ancestral recombination which is highly supported in both the Spike and Nucleocapsid gene families, from the ancestor of the *SC2-RaTG* clade (orange) to the ancestor of the *Pangolin* clade (purple). **(a)** For much of the genome, *Zhejiang* is more similar to the donor *SC2-RaTG* than the recipient *Pangolin*. **(b)** Our analysis infers HGTs from *SC2-RaTG* to *Pangolin* in the Spike and Nucleocapsid proteins with supports of 640 and 813, respectively. **(c)** In the 3’ region of the genome, *Pangolin* is often more similar to *SC2-RaTG*, especially in the Spike and parts of the Nucleocapsid gene families. However, it is difficult to clearly determine from sequence similarity alone which gene families have been affected by recombination, especially in ancestral cases such as these where the closest reference relative is the same for both donor and recipient.

### Bi-directional support may suggest a third-party donor

HGT events usually have high support for a single donor-recipient direction, but we found several HGTs that are “bi-directionally supported”, with neither strain appearing as the donor more than 60% of the time. Of the 588 inferred HGTs, 96 are bi-directionally supported, resulting in 48 pairs of strains with roughly equal support for an HGT in either direction in a given gene family. Note that the bi-directionally supported Spike HGT between *SC2-RaTG* and *Pangolin*, identified above, shows a 640-360 split and would therefore not be counted as bi-directionally supported using the conservative threshold used above.

Bi-directional HGTs may arise due to a lack of resolution in the species tree where an HGT in either direction can explain the gene tree topology equally well, as discussed in the previous case study with the Spike and Nucleocapsid HGTs between *SC2-RaTG* and *Pangolin*, or due to complex HGT scenarios where multiple HGT events occur in quick succession. The case study below demonstrates a case where support for both directions arises when the candidate HGT occurs in quick succession following another HGT from a third party. This case study also highlights a shortcoming of using primarily sequence-based approaches such as SimPlot and RDP for inferring such complex HGT scenarios. For instance, even the sophisticated RDP tool requires that the user accept or reject proposed recombinations, which inform subsequent proposals. Thus, it lacks the ability to automatically model inference uncertainty and report ambiguous cases. In contrast, our approach explicitly accounts for and reports uncertainty, which can highlight ambiguous cases for further investigation.

#### Case study: Bi-directional HGTs

We identified an HGT in the Spike gene between the strains Yunnan 2013 [YN2013] and Guizhou 2013 [Anlong-103] (Figure 6a,b). If we do not consider donor-recipient directionality, this HGT is supported in all 1000 samples. However, support is almost evenly split between each strain as the donor (503 Anlong-103, 497 YN2013). By comparing the sequence similarity of each strain to both each other and its nearest neighbor on the strain tree using SimPlot, we hypothesize that this directional ambiguity can be explained by the presence of a third strain that recombined with one of YN2013 or Anlong-103, which then recombined with the other strain.

**Fig. 6:**
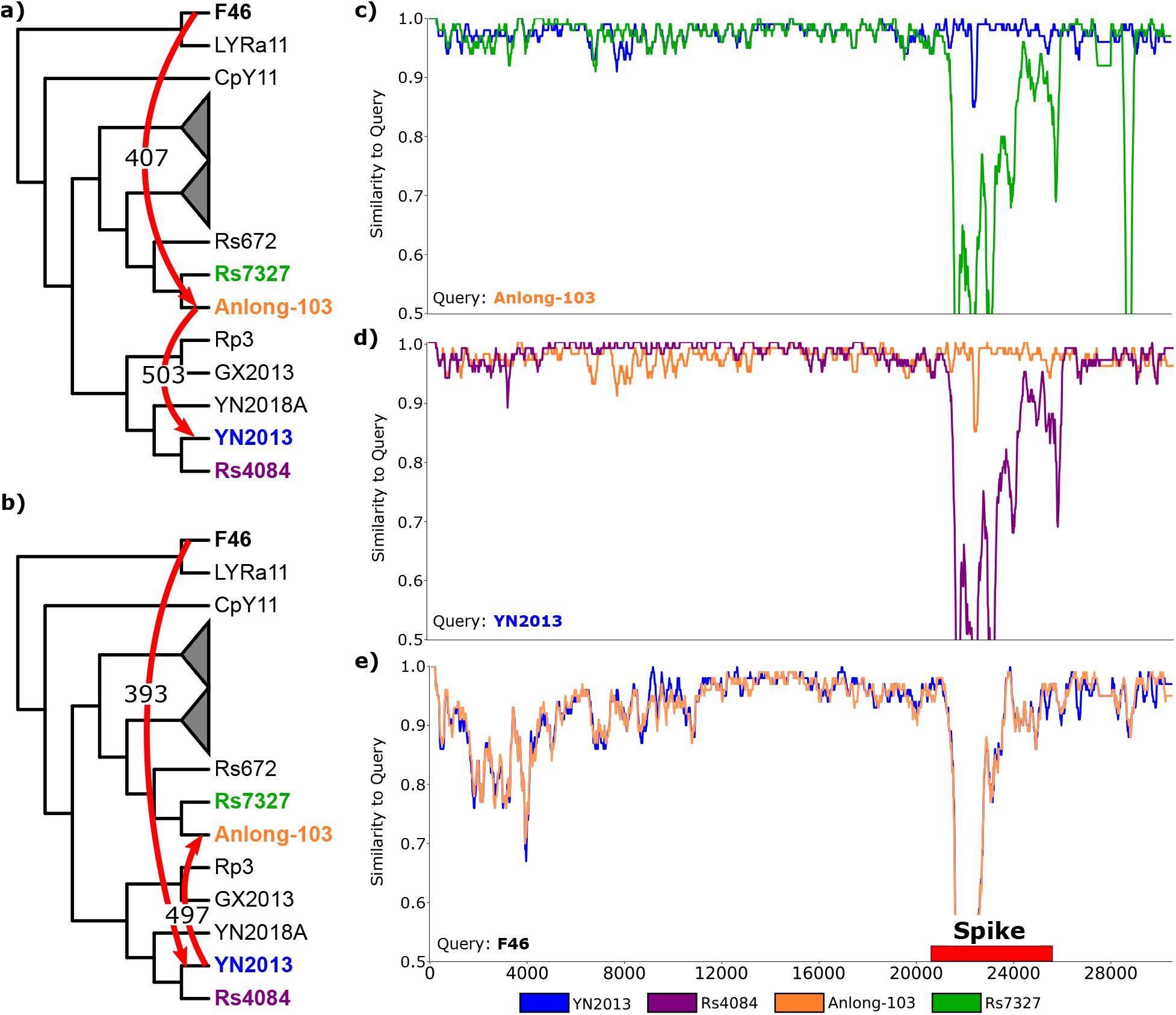
Pairs of strains with bi-directional HGTs suggest the presence of a third party donor. We present a case study of an inferred HGT between Anlong-103 (orange) and YN2013 (blue) in the Spike protein, with support of **(a)** 503 in the forward direction and **(b)** 497 in the backward direction. Such bi-directional support suggests strong evidence that an HGT occurred and a third-party was involved, but ambiguity as to the direction of the HGT. We found support for HGTs **(a)** from F46 to Anlong-103 (407 support) and **(b)** from F46 to YN2013 (393 support). **(c)** and **(d)** show Simplot analysis demonstrating both Anlong-103 and YN2013 are significantly different from their respective siblings ( RS7327, green, and Rs4084, purple) in the Spike protein. **(e)** Simplot analysis shows that both Anlong-103 and YN2013 are equally similar to a putative third-party donor F46 (black).

Using Anlong-103 as the query (Figure 6c), we found that its sibling Yunnan 2014 [Rs7327] is highly similar along the length of the genome, except in the Spike region, where there is a significant drop off in similarity. In contrast, YN2013 stays highly similar except in the variable-loop region, where there is a slight drop off in similarity. We observe the same similarity profile using YN2013 as the query with its sibling Yunnan 2012 [Rs4084] (Figure 6d). In the case of a simple unidirectional HGT event, we would expect only one of these queries to be dissimilar to its sibling.

Indeed, our HGT analysis finds third party HGTs consistent with this interpretation, with Yunnan 2012 [F46] as the third strain. We found unidirectional HGTs between Yunnan 2012 [F46] and both Anlong-103 (407 support, Figure 6a) and YN2013 (393 support, Figure 6b). SimPlot analysis supports this interpretation, with both YN2013 and Anlong-103 showing the same similarity drop-off profile to F46 through the Spike region (Figure 6e). The question of which strains recombined first is less clear, but geography and sampling times suggest that F46 and YN2013, both sampled in Yunnan province, could have recombined first followed by the recombination between YN2013 and Anlong-103 (Figure 6b).

### Recombination occurs across gene boundaries

Our reconciliation-based approach infers HGT events independently for each gene family. However, multiple such “HGTs” may result from a single recombination event across gene boundaries. To investigate this possibility, we assumed a null hypothesis that HGTs are not correlated across genes and assessed the frequency of HGT in adjacent gene families. Specifically, we assessed the *p*-value of seeing at least *t* HGTs in a window of *w* adjacent genes among the inferred donor-recipient pairs (Table S4). Here, we treat HGTs as undirected edges, with the HGT support values aggregated across both directions to account for areas of directional uncertainty. For our null distribution, we assumed a binomial distribution, where the number of trials *n* is the number of strain pairs in our data set with at least *t* HGTs. To obtain the probability of success *π*, we randomly sampled 500,000 strain pairs from our data set and independently randomly permuted the gene family ordering of each pair of strains. Out of all pairs of strains with at least *t* HGTs, we calculated *π* as the fraction of those that fit the window condition. We then computed the probability of seeing at least *k* successes, where a success corresponds to a strain pair with at least *t* HGTs in at least one window of size *w*. We investigated several combinations of (*w, t*) and rejected the null hypothesis at a significance level of *α* = 0.007 (after Bonferonni correction for 7 tests) for (*w, t*) = (2, 2), but failed to do so for larger window sizes or more HGTs. This result suggests that HGTs in adjacent pairs of gene families between two strains are likely due to a single recombination event. Thus, it should be possible to combine inferred individual HGTs with gene adjacency information to identify larger recombination regions.

## Discussion

The emergence of SARS-CoV-2 demonstrates the need to understand how novel pathogens originate by crossing species boundaries and how they adapt through recombination. In this work, we introduce virDTL, a new computational protocol for viral recombination analysis, and use it to provide a more complete picture of the evolutionary history of SARS-CoV-2 in particular and *Sarbecoviruses* in general. A key feature of virDTL is its ability to identify ancestral recombinations and provide support values for each event. virDTL leverages the Duplication-Transfer-Loss (DTL) model and accounts for multiple sources of inference uncertainty, making it a principled, model-based approach and well-suited to analyzing rapidly-evolving RNA viruses.

Many existing analyses of viral recombination often rely on comparing sequence similarity alone, using tools such as SimPlot [28] and RDP [30]. While such tools can identify recombinant strains and recombinant regions within those strains, they typically require a combinatorial exploration of query and reference sequences against which to compare the proposed recombinant and are not well suited for detecting ancestral recombinations. Their results are also hard to interpret when the strain lineages being analyzed have been affected by multiple successive recombination events. Additionally, they are unable to capture the uncertainty that arises from HGTs occurring in rapid succession in a single lineage. These tools thus work well for investigating recent recombination events in specific strains of interest, but they are difficult to use when one wants to systematically detect ancestral recombination events and precisely identify the recombining ancestral strains.

There have been two recent investigations of HGT and recombination in coronaviruses using phylogenetic reconciliation approaches [14, 29] (performed independently in parallel to the current work). Fu et al. [14] used DTL reconciliation to infer inter-host HGT events using approximately 400 coronavirus genomes, including alpha, beta, delta, and gamma coronaviruses from a variety of host species. The authors identified 5 gene clusters that were generally well-conserved among the considered genomes, used their concatenated alignments to reconstruct a coronavirus phylogeny, and reconciled it with gene trees from 20 protein families found in at least 30% of the genomes using the DTL reconciliation software RANGERDTL [4]. The resulting reconciliations were used to identify the host species that were most likely to engage in cross-host-species recombination of coronaviruses. Makarenkov et al. [29] used phylogenetic techniques to investigate patterns of HGT and recombination in 11 gene families from *Sarbecoviruses*. In particular, the authors use the HGT detection program T-Rex [6], based on bipartition dissimilarity between a strain tree and gene trees, to identify partial- and full-gene HGTs. While these investigations illustrate the potential of using phylogenetic reconciliation for studying viral evolution, neither adequately addresses key sources of HGT inference error and uncertainty, likely leading to decreased accuracy and spurious events. For instance, the analysis of Fu et al. [14] does not account for gene tree error and inference uncertainty, rooting uncertainty, and reconciliation uncertainty.

Our approach also differs significantly from that of Makarenkov et al. [29], where key differences in methodology lead to several differences in inferred events. For example, Makarenkov et al. use a single whole-genome phylogeny as their betacoronavirus strain tree, which, as our analysis suggests, has likely been affected by substantial recombination (*Methods*). While we infer transfers of the Spike and Nucleocapsid genes between the SC2-RaTG clade and the Guangxi pangolin clade, Makarenkov et al. instead infer transfers between RaTG13 to Guangxi pangolin. While our analyses differ by one branch, Makarenkov et al. only infer a transfer because they include Guangdong pangolins. That is, their analysis might not have identified a transfer using our set of species, which highlights the potential increased sensitivity of our approach. Makarenkov et al. also do not infer transfers of the Nucleocapsid gene between SC2-Zhejiang and more distant relatives. This discrepancy is likely due to the differences in species tree topology, where Makarenkov et al. place the Zhiejiang clade as the outgroup of the SC2-RaTG-pangolin lineage. However, based on sequence similarity, it is likely that the Zhiejiang clade is a sister clade to SC2-RaTG clade as suggested by the NRR-B species tree. At the same time, Makarenkov et al. infer several gene transfers that we do not find in our analysis. For example, they postulate partial gene transfers of ORF1ab and Membrane genes between the Zhejiang clade and SC2-RaTG clade and complete gene transfers of ORF3a, ORF8, and ORF10 between the ancestor of Guangdong pangolins and Wuhan-Hu-1 and the Zhejiang clade. These events would likely not occur using a non-recombinant strain tree such as our NRR-B tree, which places the Zhejiang clade as a sister clade to SC2-RaTG. In addition, while Makarenkov et al. do implicitly consider gene tree inference uncertainty, by considering bootstrap values along gene tree edges to assign support values for inferred HGT events, they do not perform gene tree error-correction, which has been shown to result in significant improvements in downstream HGT inference accuracy [5, 20, 39]. Lastly, Makarenkov et al. also use an older HGT detection tool, T-Rex [6], which is not based on DTL reconciliation and does not explicitly address HGT inference uncertainty due to multiple optima.

Our analysis of *Sarbecovirus* evolutionary history lends additional support to the growing body of work that suggests horseshoe bats as the most recent zoonotic origin of the SARS-CoV-2 lineage. Similarity of the ribosome binding domain (RBD) of the Spike protein between SARS-CoV-2 and several pangolin strains has led to the hypothesis of an intermediate pangolin reservoir for the virus [25, 50]. However, our analysis suggests that this similarity is due to a recombination event between the ancestor of SARS-CoV-2 [Wuhan-Hu-1] and RaTG13 and the ancestor of the Guangxi pangolins. Sequence evolution through mutations or recombination with an unsampled strain would account for the divergence of RaTG13 in this region, consistent with the observations of Boni et al. [7].

Our approach has several limitations that are worth noting. Most importantly, we analyze each gene family separately and thus cannot infer recombination events that affect only parts of genes. Moreover, uncertainly and error in HGT inference and in assigning donors and recipients can make it difficult to infer larger recombination events that affect multiple genes. This limitation can be partially addressed by using a window-based analysis, rather than a gene-based analysis, but small windows risk having too little meaningful phylogenetic signal while large windows risk averaging over several different overlapping recombination events. Another limitation of our approach and analysis is that it ignores low-support HGTs. Low-support HGTs cannot be disregarded altogether, especially when the strains being analyzed contain short genes. Short genes, such as the E, ORF7b, and ORF10 gene families in our *Sarbecovirus* analysis, often have less phylogenetic signal and thus more uncertain gene tree topologies and inferred events. A closer analysis of low-support HGTs, especially those affecting short genes, may thus lead to additional evolutionary insights.

## Acknowledgements

The authors would like to thank Irwin Jungreis and Rachel Sealfon for help with dataset assembly, and Dong-Hun Lee for helpful discussions on coronavirus evolution.

## Funding

SZ and MSB were supported in part by NSF grant IIS-1553421. SS and BB were supported in part by NIH grants R01-GM081871 and R35-GM141861. YW was supported by NSF grant IIS-1751399.

## Methods

### *Sarbecovirus* strain selection

For our analysis we selected 54 strains from the *Sarbecovirus* subgenus of the *Betacoronavirus* genus, with 42 strains from bats, 5 from pangolins, 5 from civet cats, and 2 from humans. We limited our strain selection to only the *Sarbecovirus* subgenus since genomes outside this subgenus, such as MERS-CoV, are generally too divergent from SARS-CoV-2 [21], and analyses including such distant strains can fail to cleanly identify gene families or can result in phylogenetic artifacts such as long branch attraction. For example, even the closest relative outside the *Sarbecovirus* subgenus, Hibecovirus Bat Hp-betacoronavirus/Zhejiang2013, shows no detectable homology across ORFs 6, 7a, 7b, and 8 [21].

Our choice of the 54 *Sarbecovirus* strains was driven by two considerations. First, we wanted to sample broadly from available *Sarbecovirus* whole genomes in order to adequately capture strain diversity, and second, we wanted to include strains with potential relevance for SARS-CoV-2 evolution. Accordingly, we started with the collection of 44 broadly sampled *Sarbecovirus* strains (one SARS-CoV-2 strain, one SARS-CoV strain, and 42 strains from bats) used in Jungreis et al. [21], and augmented that collection with strains hosted in civet cats and pangolins due to their proposed role as zoonotic origins for the SARS (2003) [18] and SARS-CoV-2 (2019) [50] pandemics. A complete list of the chosen strains is available in Supplementary Table S1. For each strain, the complete genome sequence was obtained from the NCBI sequence database [32].

Throughout this study, we reference results of Boni et al. [7] and Makarenkov et al. [29]. Boni et al. include 19 strains in their analysis that we leave out, including 279 2005, JL2012, JTMC15, SX2013, Rs4874, RsSHC014, Rs3367, Longquan 140, HKU3-[2–6,8–11,13], and Pangolin-CoV. As these strains have a close relative HKU-3-[1,7,12] that is included in our analysis, and the strains do not represent new hosts, their exclusion should not alter results substantially. We include two additional strains, 16BO133 and 273 2005. Makarenkov et al. includes 6 strains that we leave out, including Guangdong Pangolin 1 2019, Guangdong Pangolin P2S 2019, HKU3-6, and three SARS-CoV-2 strains (Australia VIC231 2020, USA UT 00346 2020, Hu Italy TE4836 2020). As mentioned in Makarenkov et al., the SARS-CoV-2 strains are very similar and therefore do not add further information to the analysis.

### Strain tree reconstruction and dating

Given the importance of strain tree accuracy on the accuracy of HGT inference through phylogenetic reconciliation, we investigated three candidate genomic regions to reconstruct a dated strain tree. As a baseline, we constructed a whole genome (*WG*) strain tree based on a whole genome alignment of the 54 genomes. However, coronaviruses are highly recombinant, so it is preferable to use a generally non-recombining, or minimally recombining, genomic region to accurately capture the divergence of viral lineages. We therefore selected two putative non-recombinant regions (*NRR-B* [4,000-9,000 base pairs] and *NRR-A* [13,000-18,000 base pairs]) previously identified by Boni et al. [7].

For each region/whole genome, we aligned the 54 sequences using Muscle v.3.8.31 [12] and estimated a dated strain tree using BEAST v.1.10.4 [42]. Following Boni et al. [7], we used a GTR+*Γ* substitution model and an uncorrelated relaxed clock model with a log-normal distribution. We used a normal distribution with mean 0.00078 and standard deviation 0.0003 as an informative rate prior, based on estimated rates for MERS-CoV [7], and ran BEAST until chains were sufficiently mixed, generally for more than 10 million iterations, with effective sample sizes greater than 100 for branch lengths and root ages. We rooted each strain tree using the outgroup containing the strains from Bulgaria 2008 [BM48-31] and Kenya 2007 [BtKY72], which were identified in Boni et al. [7] as the most evolutionarily distant strains. Given the rooted strain trees, we again ran BEAST until chains were sufficiently mixed, using topology-preserving operations only to estimate divergence times for each ancestral strain.

The three *Sarbecovirus* strain trees, corresponding to NRR-A, NRR-B, and whole genome, each had distinct topologies (Figure 2). To assess the magnitude of topological divergence between these trees, we computed the normalized unrooted Robinson-Foulds distance between them [36] (Table 1, top rows). Though several important clades appear largely conserved across the three trees (Figure 2a), the three trees are highly divergent (RF: 0.62–0.80), suggesting that the *Sarbecovirus* subgenus is influenced by substantial recombination. This result in turn implies that the WG tree should not be directly used as the strain tree, and motivates the need for constructing a reliable strain tree using a non-recombinant (or minimally recombinant) region. We found a specific instance of recombination that occurred ancestral to SARS-CoV-2 [Wuhan-Hu-1] to be of particular interest, as it explains a key difference in topology between the trees inferred using NRR-A and NRR-B. Specifically, viral strains CoVZC45 and CoVZXC21 (*Zhejiang clade*) are placed in a different location in each of the three strain trees (Figure 2a). In the WG strain tree, the clade containing Wuhan-Hu-1 and RaTG13 (*SC2-RaTG clade*) and the clade containing the *Pangolin* viral strains (*Pangolin clade*) are most closely related, with the Zhejiang clade as the next closest relative. In the NRR-B strain tree, the Zhejiang and SC2-RaTG clades are sisters, with the Pangolin clade as the next closest relative, suggesting a recombination somewhere outside of NRR-B. Finally, in the NRR-A strain tree, the Zhejiang clade does not group with either the SC2-RaTG or Pangolin clades, and the two strains instead group with 3 viral strains from Hong Kong (*HKU clade*).

**Table 1:** Three candidate strain trees are highly divergent. (*Top*) The relatively high Robinson-Foulds (RF) distances between pairs of strain trees indicates substantial differences between the tree topologies, and suggests that substantial recombination has occurred throughout the *Sarbecovirus* subgenus. This result motivates the need for constructing a reliable strain tree using a non-recombinant region. (*Bottom*) We report the average normalized RF distance between trees constructed within a genomic region. We divided the genome into 1000-base pair windows with a 500-base pair offset, reconstructed a phylogeny on this window using RAxML, and computed all pairwise RF distances between the windows. We show the average internal pairwise RF distance for 5000-base pair windows and compare to the average internal pairwise RF distance for the two putative non-recombinant regions. **NRR-B is more internally consistent than other genomic regions**, which suggests less recombination in this region and motivates its use as our strain tree for reconciliation.

| Genome Region | Normalized RF Distance |
| --- | --- |
| Whole Genome vs. NRR-B | 0.660 |
| Whole Genome vs. NRR-A | 0.620 |
| NRR-A vs. NRR-B | 0.800 |
| 0 - 5000 | 0.522 |
| **4000 - 9000 (NRR-B)** | **0.496** |
| 5000 - 10000 | 0.519 |
| 10000 - 15000 | 0.559 |
| 13000 - 18000 (NRR-A) | 0.607 |
| 15000 - 20000 | 0.586 |
| 20000 - 25000 | 0.549 |
| 25000 - 30000 | 0.561 |

#### Recombination within NRR-A and NRR-B

To assess whether recombination might affect the NRR-A and NRR-B trees, we constructed phylogenies for 1000-base pair windows with a 500-base pair offset along the entire length of the genome. We then computed the average pairwise Robinson-Foulds distance between all trees in a 5000-base pair window, and similarly computed the average internal pairwise distance between trees in each non-recombinant region (Table 1, bottom rows). We find that average RF distance within NRR-B (0.496) is smaller than NRR-A (0.607) and also smaller than other 5000-base pair windows along the length of the genome (0.519–0.586). A higher average internal RF distance indicates increased phylogenetic incongruency between windows within each region, which suggests recombination may have occurred within the region.

#### Recombination across NRR-A

We performed further analysis to determine if the discrepancy in NRR-A and NRR-B strain tree topologies is a result of recombination in the putative NRR-A. Specifically, using the NRR-B tree as our viral strain tree, we found evidence for an ancestral HGT between the ancestors of the Zhejiang and HKU clades (Figure 2d). This HGT is further supported by sequence similarity. Using SimPlot [28], we compared a query of Zhejiang 2017 [CoVZC45] against SARS-CoV-2 [Wuhan-Hu-1], Yunnan 2013 [RaTG13] and the three Hong Kong strains HKU3 1, HKU3 7, and HKU3 12. While Zhejiang 2017 is most similar to the SC2-RaTG clade for most of the genome, it is more similar to the HKU clade between 11,857 and 20,677 base pairs, which contains NRR-A (Figure 2c). We note that this HGT was not inferred using the MAD-rooted gene tree. Nonetheless, the similarity between the Zhejiang and HKU clades in this region indicates that recombination has in fact occurred in NRR-A, making it unsuitable to construct a strain tree using this part of the genome. This finding is consistent with the conclusions of Boni et al. [7], where they note that the Zhejiang clade needed to be removed to maintain a clean non-recombinant signal in this region.

Given that (i) the whole genome tree is generally unreliable as a strain tree for reconciliation analysis due to widespread recombination across the genome, (ii) NRR-A is far less internally consistent than NRR-B and, (iii) a major topological discrepancy in the NRR-A tree is likely the result of an ancestral recombination, we used the NRR-B tree as our viral strain tree for the remainder of our analyses.

### Gene tree reconstruction, error-correction, and rooting

We constructed gene trees for each of 11 gene families (Figure 2b). Most strains in our dataset were already annotated with genes from all 11 gene families. For each annotated gene, we extracted and used the longest protein sequence for that gene. For strains that did not have all 11 genes annotated, we aligned their full genome to the longest annotated gene sequence among other strains and extracted the overlapping alignment. Almost all gene families, with the exception of ORF1ab, Spike, and Nucleocapsid, were unannotated in at least one strain. Using this approach, we were able to confidently identify the missing genes for an additional 5 gene families, resulting in a total of 8 complete gene families (with one gene from each of these gene families present in each strain). Two of the gene families, ORF10 and ORF6, could not be detected in strains KJ473815.1 and KJ473816.1. Finally, ORF8, which was initially annotated in only 31 of the 54 strains, could not be detected in 10 strains. For each gene family, we aligned nucleotide gene sequences using Muscle v.3.8.31 [12] and then reconstructed gene trees using RAxML v.8.2.11 [40] using 100 fast bootstrap replicates under a GTR+*Γ* substitution model.

Since there is often considerable uncertainty and error in reconstructed gene trees, generally due to insufficient phylogenetic signal, we error-corrected each RAxML gene tree using TreeFix-DTL [5] with default parameters. TreeFix-DTL aims to find a “statistically equivalent” gene tree topology that minimizes the DTL reconciliation cost against a given species/strain tree, and has been shown to be highly effective in error-correcting gene trees, leading to a substantial reduction in the number of false positive HGTs inferred through DTL reconciliation [5]. To further account for uncertainty in inferring gene tree topologies, we applied TreeFix-DTL 10 times to each RAxML gene tree and used all 10 error-corrected gene trees for each gene family in our analysis.

Most phylogenetic reconstruction methods, including RAxML and TreeFix-DTL, yield unrooted trees that can often be difficult to root accurately. For example, even the root of the SARS-CoV-2 phylogeny remains uncertain, with different rooting methods yielding different root positions [35]. Thus, to account for uncertainty in gene tree rooting, we rooted each error-corrected gene tree using two different methods, OptRoot [4], which seeks a rooting that minimizes the DTL reconciliation cost between the gene tree and species/strain tree, and Minimum Ancestor Deviation (MAD) rooting [47], which roots the gene tree at the edge that minimizes the mean relative deviation from the molecular clock. These two rooting methods have been shown to be among the most accurate for prokaryotic gene families [48]. By default, we report results based on OptRoot-rooted gene trees, but all HGTs are supported by MAD-rooted gene trees unless otherwise stated.

### Reconciliation analysis and accounting for HGT inference uncertainty

We reconciled each of the rooted, error-corrected gene trees (10 per gene family) to the NRR-B strain tree using RANGER-DTL 2.0 [4] with default parameters. Since there often exist multiple equally optimal DTL reconciliations of a given gene tree and strain tree [3], we uniformly random sampled 100 optimal reconciliations (per rooting) for each pair of gene and strain trees. Such uniform random sampling makes it possible to assign a support value to each inferred HGT event based on how frequently that event is inferred among all optimal DTL reconciliations. These support values can then be used to distinguish between HGTs that are well-supported by DTL reconciliation, despite multiple optima, and those that are not.

As expected, we observed that the computed reconciliations invoked a negligible number of gene duplication events. Specifically, for 8 of the 11 gene families, no gene duplications were invoked in any of the 1000 reconciliation samples for each gene family. For the ORF1ab, Envelope, and Nucleocapsid gene families, we observed 0.21, 0.1, and 0.02 gene duplications, respectively, on average, in the 1000 reconciliation samples. This result implies that almost all topological inconsistencies between the gene trees and the strain tree can be explained using HGT events, which is consistent with the assumption that, in viruses, any topological discordance between the gene trees and strain tree is likely a result of recombination.

### Recombination across gene boundaries

Though we primarily focused our analysis at the level of individual gene families, we performed statistical tests to investigate whether our results can be used to detect recombination of larger genomic fragments (e.g., multiple genes) in the *Sarbecovirus* subgenus. In particular, we investigated how often two or more contiguous genes are likely to be transferred between the same donor-recipient pair. We also used statistical analysis to assess if such “multi-gene” transfers were observed at a higher rate than would be expected by chance if recombination events affected only one gene at a time (*Results*).

## Supplementary Material

**Fig. S1:**
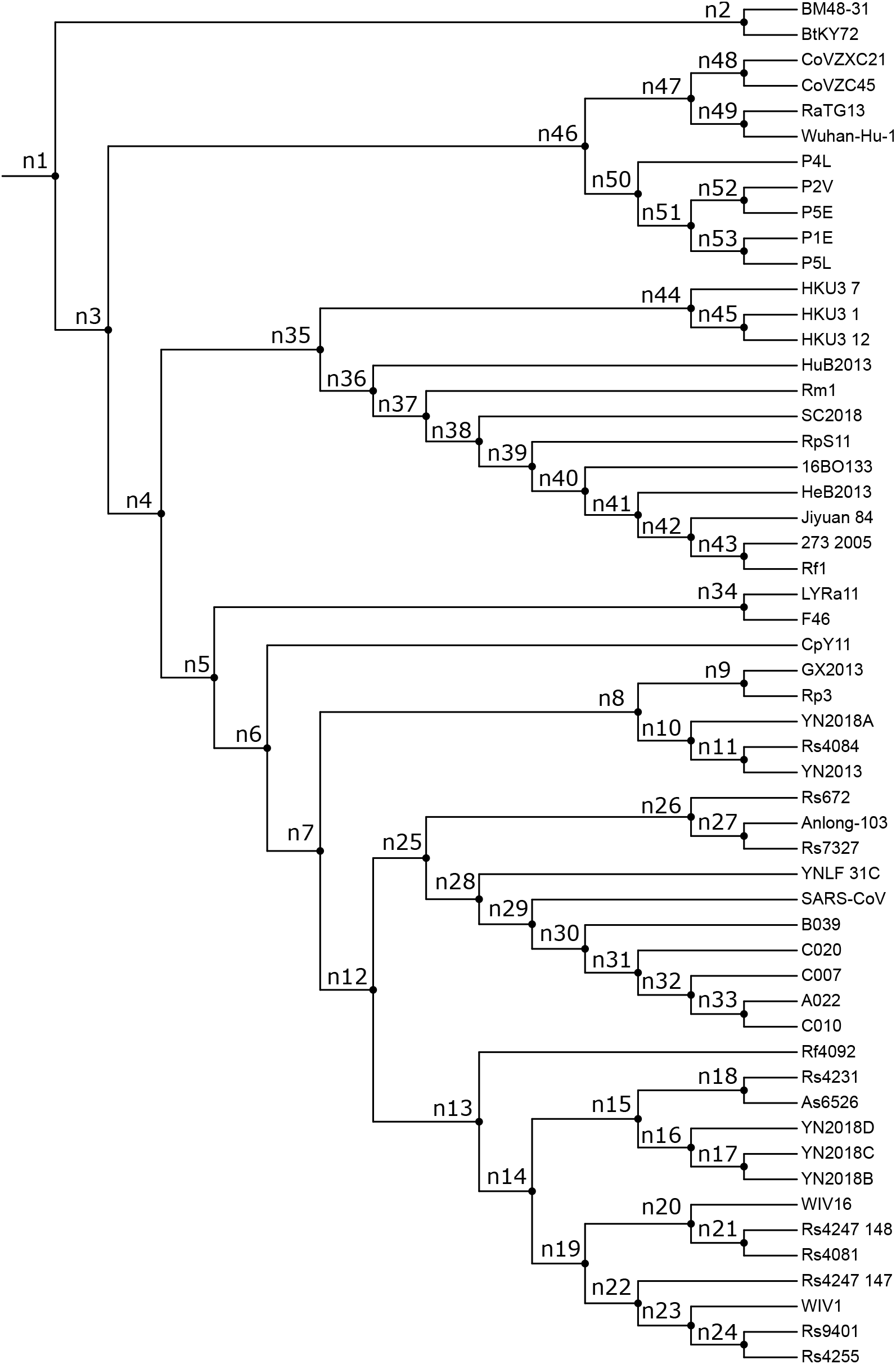
Full strain tree (NRR-B) and internal node labels.

**Fig. S2:**
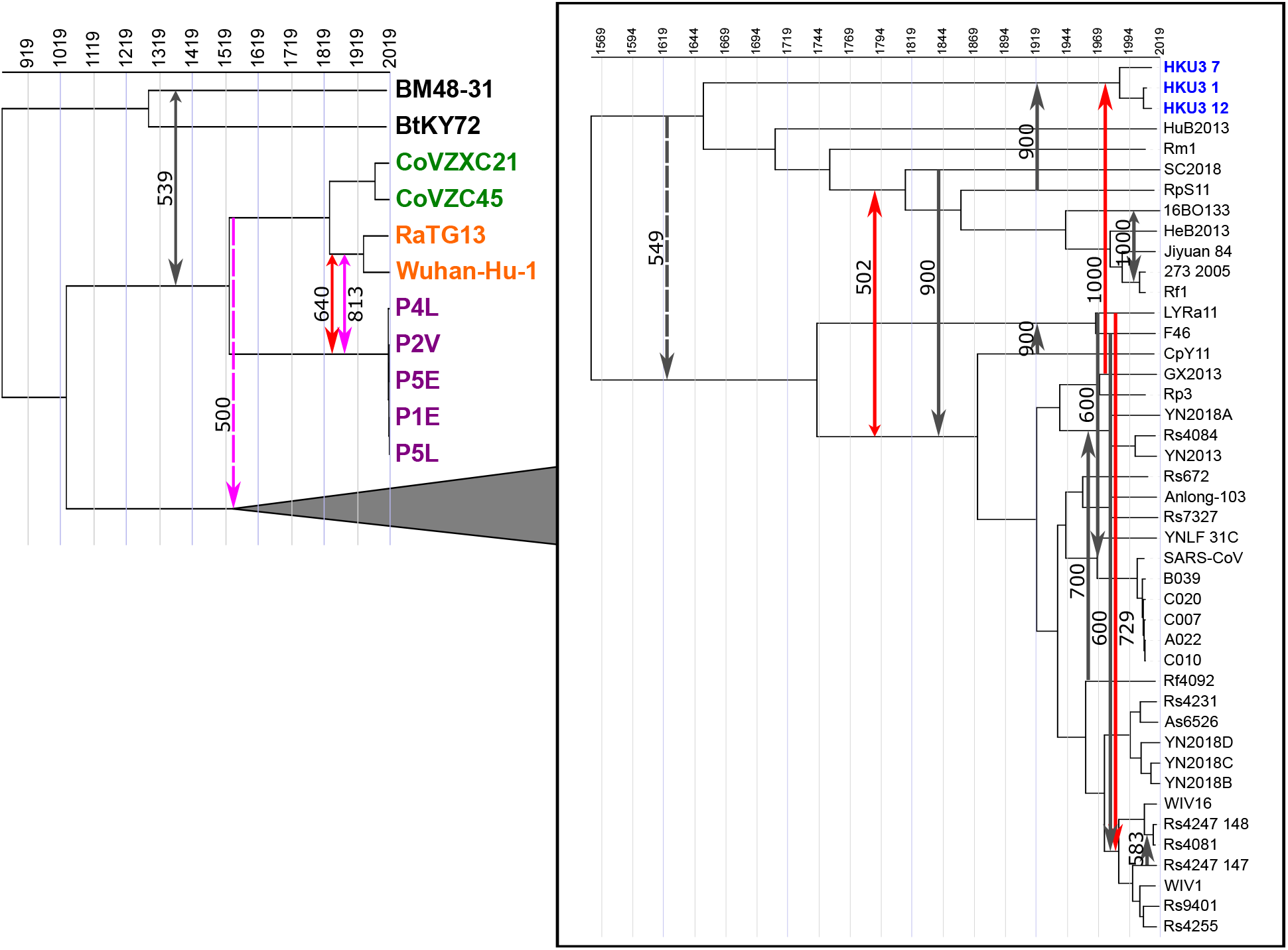
Highly-supported time-consistent HGTs in the *Sarbecovirus* subgenus. Time-consistent HGTs with an ancestral recipient and greater than 500 support are shown on a dated strain tree. Support values are shown for OptRoot-rooted gene trees, with transfers in the Spike gene (red) and Nucleocapsid gene (pink) highlighted. Smaller arrows indicate there also exists an HGT with at least 100 support in the reverse direction, suggesting directional uncertainty. All HGTs shown are also supported using MAD-rooted gene trees except one transfer in the Nucleocapsid gene and one in the Membrane gene (dashed lines).

**Fig. S3:**
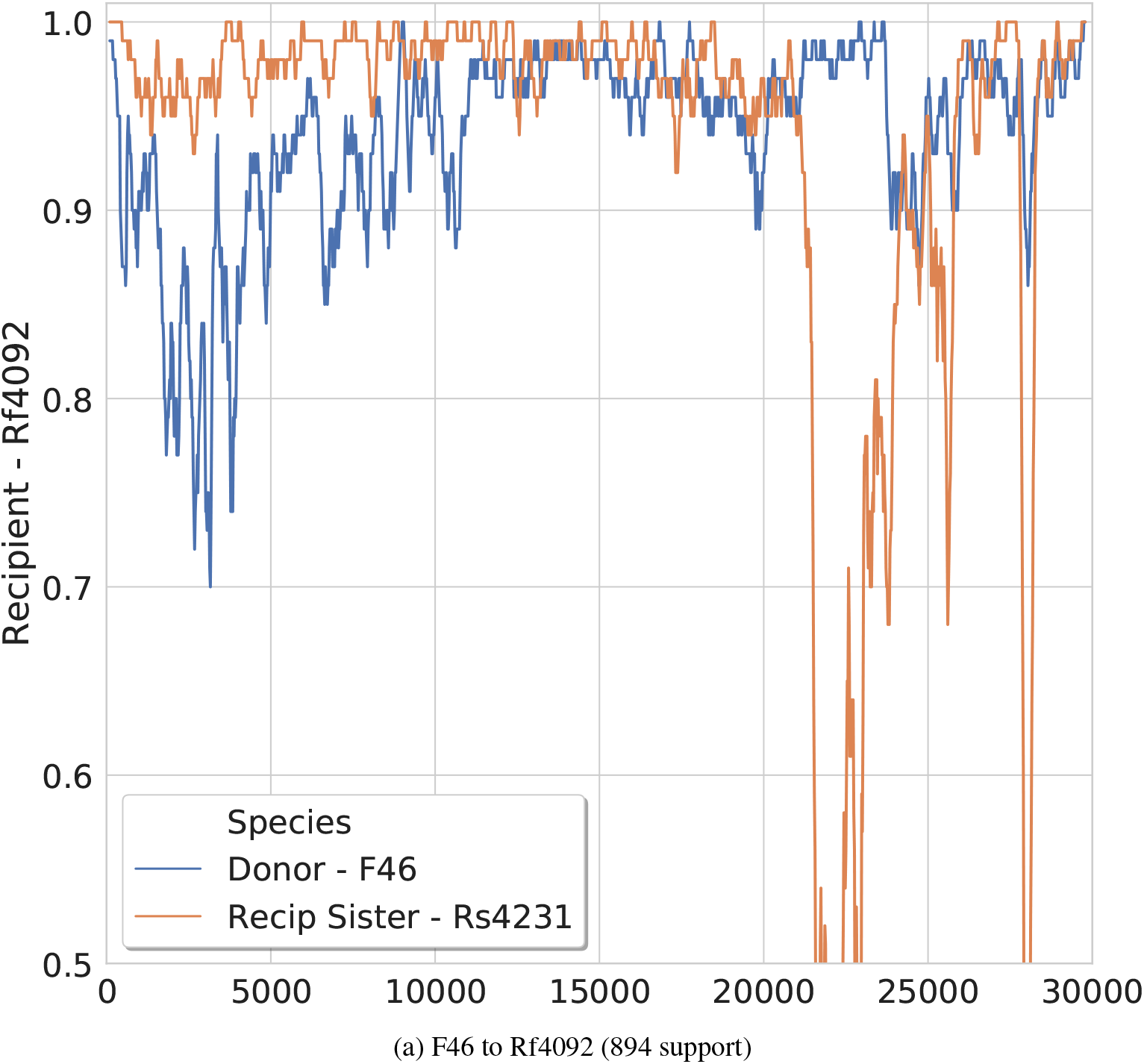
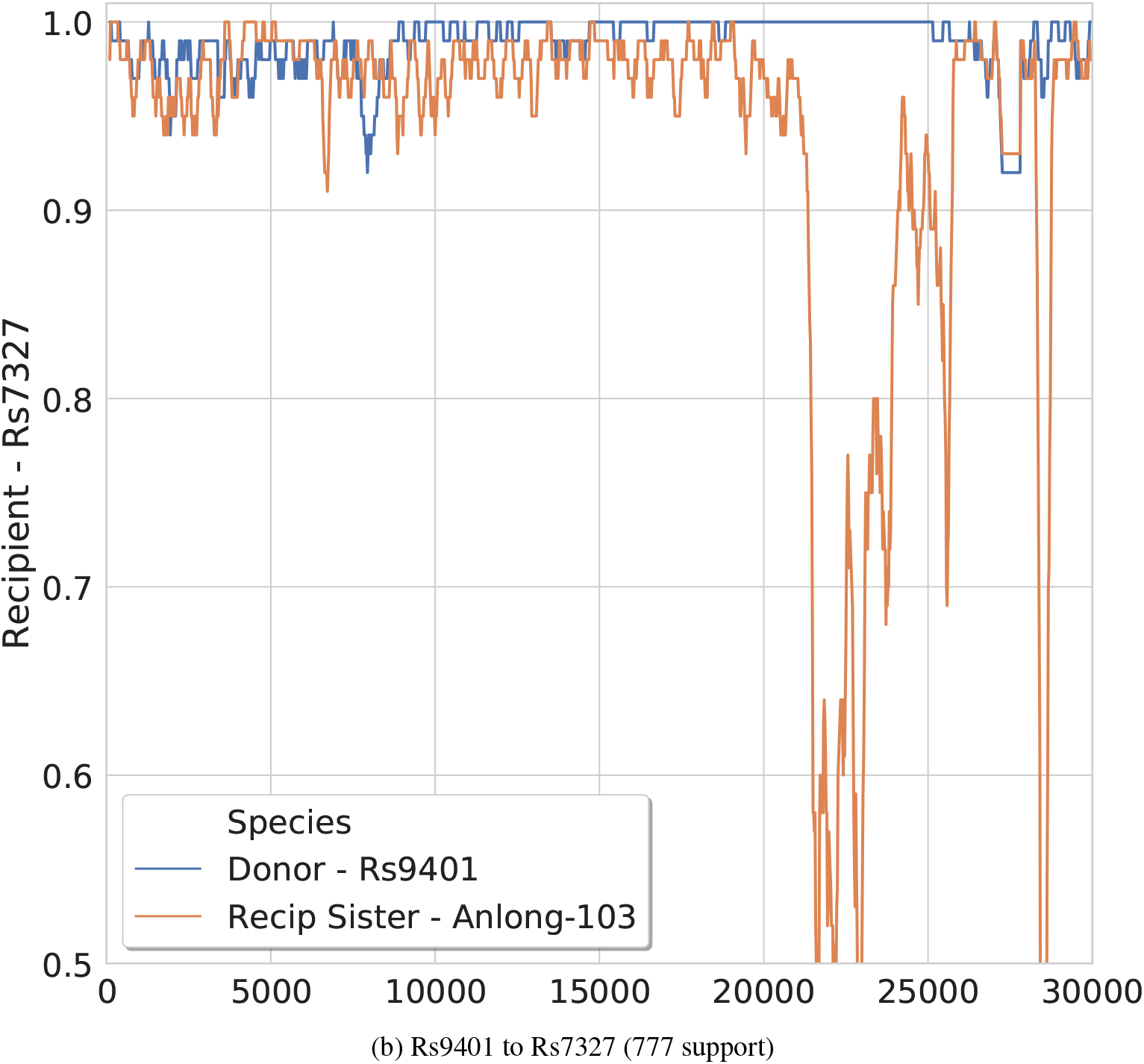
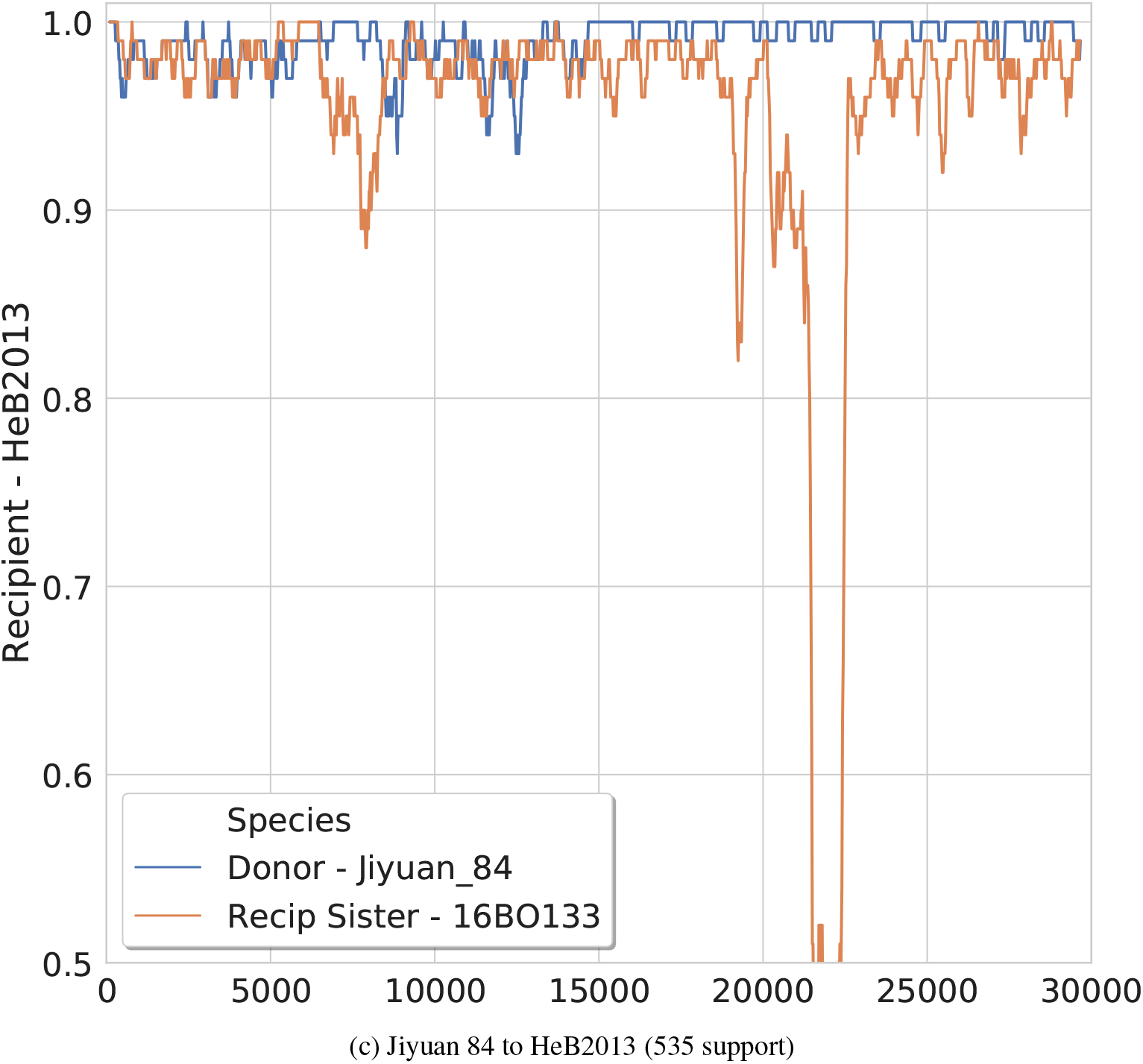
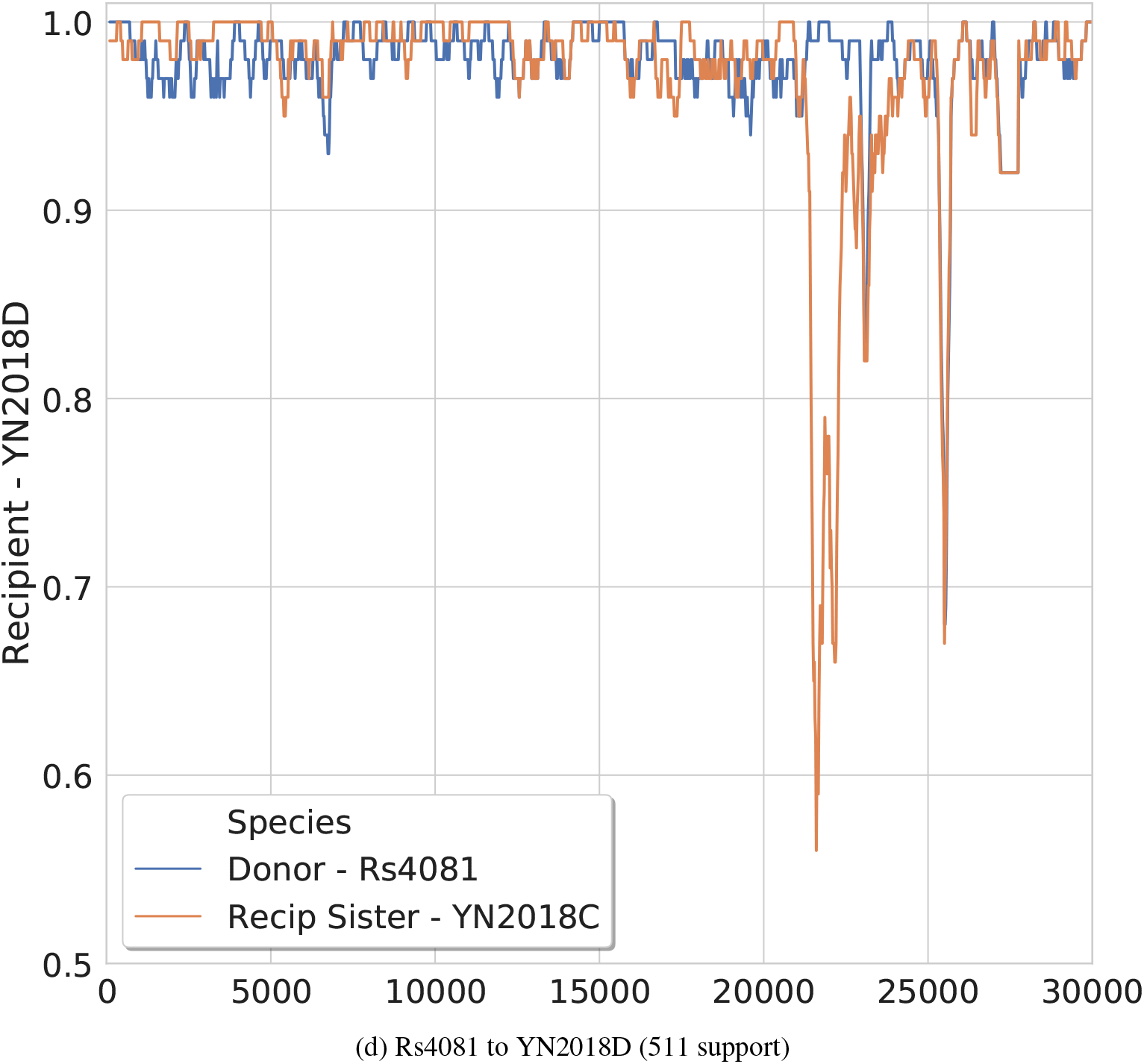
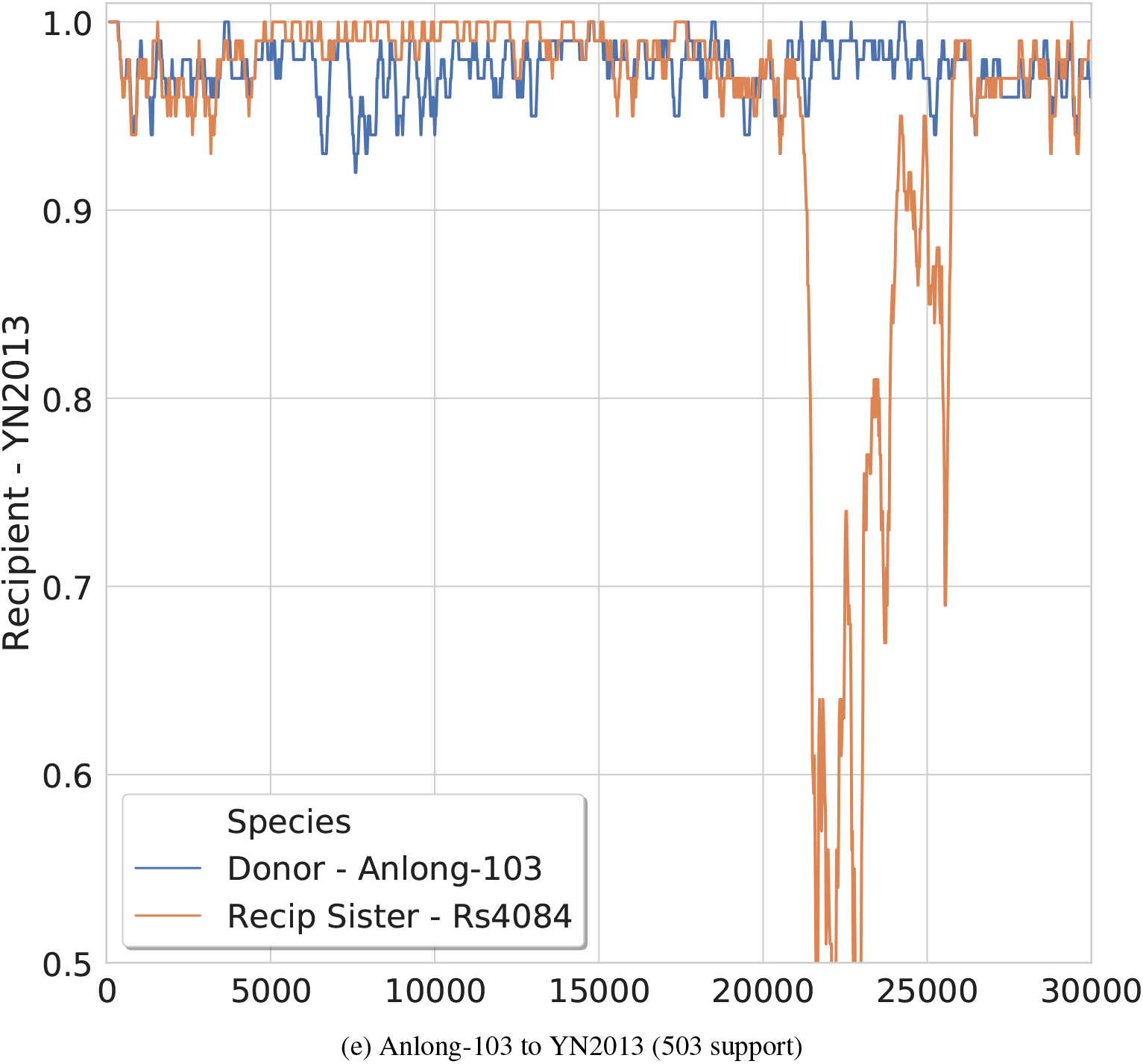
SimPlot Validation of leaf-to-leaf HGTs. Part (a): SimPlot for highly supported leaf-to-leaf HGT in the Spike gene family from F46 to Rf4092. Figure continued on next page. Part (b): SimPlot for highly supported leaf-to-leaf HGT in the Spike gene family from Rs9401 to Rs7327. Figure continued on next page. Part (c): SimPlot for highly supported leaf-to-leaf HGT in the Spike gene family from Jiyuan 84 to HeB2013. Figure continued on next page. Part (d): SimPlot for highly supported leaf-to-leaf HGT in the Spike gene family from Rs4081 to YN2018D. Figure continued on next page. Part (e): SimPlot for highly supported leaf-to-leaf HGT in the Spike gene family from Anlong-103 to YN2013.

**Table S1:**
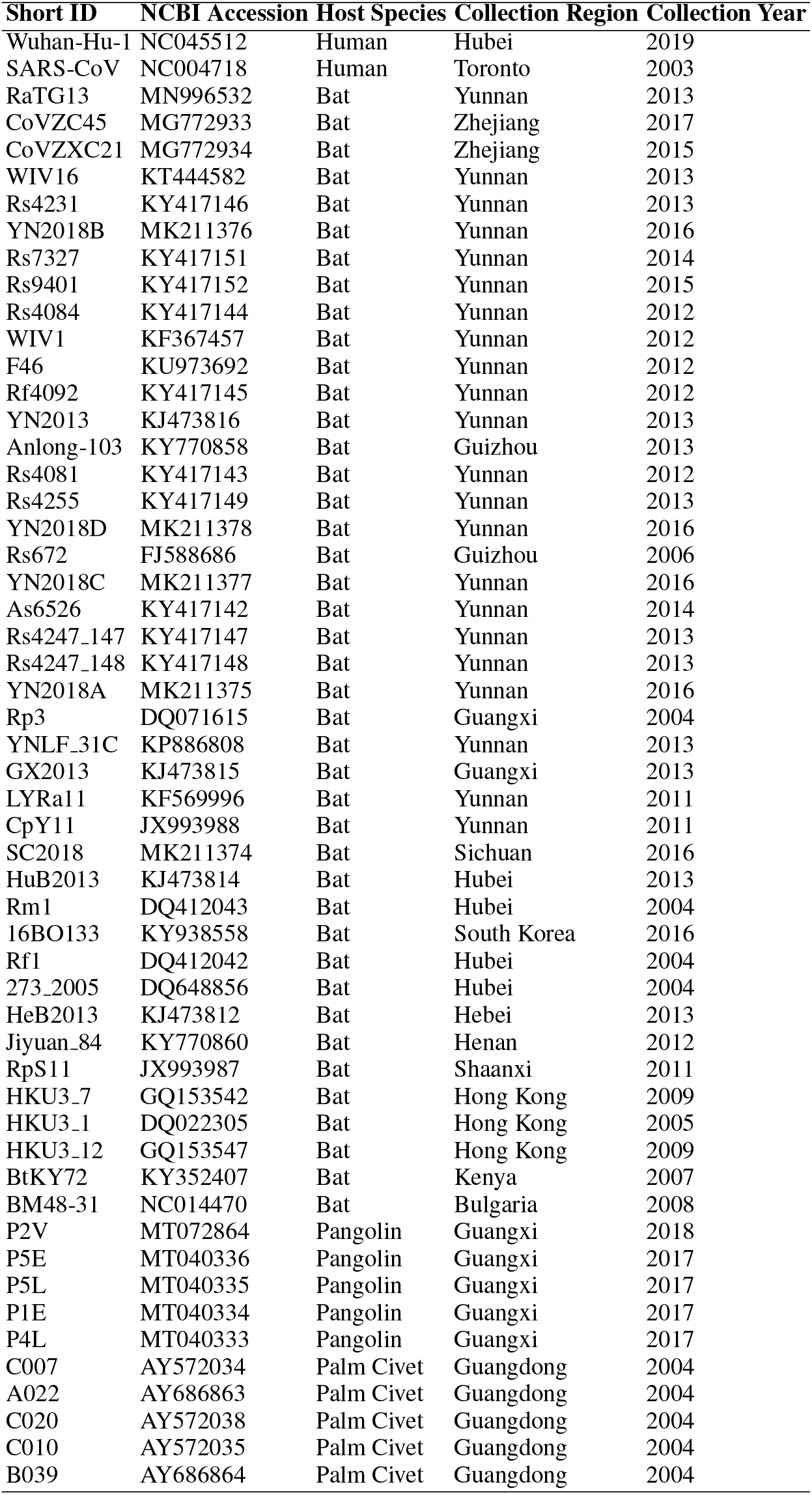
List of strains included in analysis.

**Table S2:** All HGTs with ≥ 100 support found by our analysis. See separate spreadsheet.

**Table S3:** Number of HGTs per gene family. For each gene family, we show the alignment length and the number of HGT events found at varying levels of support: at least 100 (95^th^ percentile), at least 500 (99^th^ percentile), and at least 808 (99.5^th^ percentile).

| Gene Family | Alignment Length<br>(base pairs) | HGTs |  |  |
| --- | --- | --- | --- | --- |
| | | $\geq 100$ support | $\geq 500$ support | $\geq 808$ support |
| ORF1ab | 21,465 | 42 | 11 | 3 |
| S | 3,896 | 54 | 13 | 4 |
| ORF3a | 836 | 47 | 10 | 6 |
| E | 231 | 51 | 0 | 0 |
| M | 687 | 63 | 11 | 3 |
| ORF6 | 186 | 50 | 6 | 2 |
| ORF7a | 376 | 65 | 12 | 4 |
| ORF7b | 135 | 48 | 0 | 0 |
| ORF8 | 389 | 68 | 4 | 1 |
| N | 1,271 | 74 | 9 | 2 |
| ORF10 | 117 | 26 | 2 | 0 |
| Total |  | 588 | 78 | 25 |

**Table S4:** Inferred HGTs in adjacent gene families likely recombined in a single event. We randomly sampled 500,000 strain pairs from our data and randomly permuted the gene family ordering to estimate the probability *π* that a pair of strains with at least *t* supported HGTs has at least one window of size *w* with *t* HGTs in it by random association. We then performed a one-sided binomial test with probability of success *π*, *k* strain pairs in our data that fit the (*w, t*) window condition, and *n* strain pairs that have at least *t* supported HGTs. Bold text indicates significance at *α* = 0.007, after Bonferroni correction for 7 hypotheses tested. We find that when HGTs are inferred between the same pair of strains for two adjacent gene families, they were likely transferred together, but fail to make similar claims for larger numbers of grouped genes.

| $w$ | $t$ | $\pi$ | $n$ | $k$ | $p$ |
| --- | --- | --- | --- | --- | --- |
| 2 | 2 | 0.2906 | 39 | 85 | **0.0007** |
| 3 | 2 | 0.4810 | 44 | 85 | 0.2848 |
| 3 | 3 | 0.1115 | 5 | 23 | 0.1053 |
| 4 | 3 | 0.2582 | 9 | 23 | 0.1136 |
| 4 | 4 | 0.0507 | 2 | 6 | 0.0336 |
| 5 | 4 | 0.1637 | 3 | 6 | 0.0595 |
| 5 | 5 | 0.0258 | 1 | 2 | 0.0509 |

**Table S5:** Nodes with top 5% of HGTs in subtree rooted at node (normalized for size of subtree).

| Node | Node In | Node Out | Tree In (norm) | Tree Out (norm) |
| --- | --- | --- | --- | --- |
| Rs4084 | 14 | 26 | 14.00 | 26.00 |
| n26 | 9 | 10 | 16.33 | 17.67 |
| n18 | 4 | 10 | 16.00 | 17.50 |
| Rs7327 | 15 | 17 | 15.00 | 17.00 |
| n34 | 8 | 3 | 12.50 | 19.00 |
| SC2018 | 20 | 11 | 20.00 | 11.00 |
| n24 | 5 | 10 | 17.00 | 14.00 |

## Notes

https://github.com/suz11001/virDTL

### Competing Interest Statement

The authors have declared no competing interest.

## References

1. K. G. Andersen, A. Rambaut, W. I. Lipkin, E. C. Holmes, and R. F. Garry. The proximal origin of sars-cov-2. Nature Medicine, 26(4):450–452, Apr. 2020.

2. M. S. Bansal, E. J. Alm, and M. Kellis. Efficient algorithms for the reconciliation problem with gene duplication, horizontal transfer and loss. Bioinformatics, 28(12):283–291, 2012.

3. M. S. Bansal, E. J. Alm, and M. Kellis. Reconciliation revisited: Handling multiple optima when reconciling with duplication, transfer, and loss. Journal of Computational Biology, 20(10):738–754, 2013.

4. M. S. Bansal, M. Kellis, M. Kordi, and S. Kundu. RANGER-DTL 2.0: rigorous reconstruction of gene-family evolution by duplication, transfer and loss. Bioinformatics, 34(18):3214–3216, 2018.

5. M. S. Bansal, Y.-C. Wu, E. J. Alm, and M. Kellis. Improved gene tree error correction in the presence of horizontal gene transfer. Bioinformatics, 31(8):1211–1218, 2015.

6. A. Boc, A. B. Diallo, and V. Makarenkov. T-REX: a web server for inferring, validating and visualizing phylogenetic trees and networks. Nucleic Acids Research, 40(W1):W573–W579, 06 2012.

7. M. F. Boni, P. Lemey, X. Jiang, T. T.-Y. Lam, B. W. Perry, T. A. Castoe, A. Rambaut, and D. L. Robertson. Evolutionary origins of the sars-cov-2 sarbecovirus lineage responsible for the covid-19 pandemic. Nature Microbiology, 5(11):1408–1417, Nov. 2020.

8. Z.-Z. Chen, F. Deng, and L. Wang. Simultaneous identification of duplications, losses, and lateral gene transfers. IEEE/ACM Trans. Comput. Biology Bioinform., 9(5):1515–1528, 2012.

9. L. A. David and E. J. Alm. Rapid evolutionary innovation during an archaean genetic expansion. Nature, 469:93–96, 2011.

10. A. Davin, E. Tannier, T. Williams, et al. Gene transfers can date the tree of life. Nature Ecology and Evolution, 2:904–909, 2018.

11. J.-P. Doyon, C. Scornavacca, K. Y. Gorbunov, G. J. Szöllosi, V. Ranwez, and V. Berry. An efficient algorithm for gene/species trees parsimonious reconciliation with losses, duplications and transfers. In E. Tannier, editor, RECOMB-CG, volume 6398 of Lecture Notes in Computer Science, pages 93–108. Springer, 2010.

12. R. C. Edgar. MUSCLE: multiple sequence alignment with high accuracy and high throughput. Nucleic Acids Research, 32(5):1792–1797, Mar. 2004.

13. D. Forni, R. Cagliani, M. Clerici, and M. Sironi. Molecular evolution of human coronavirus genomes. Trends in Microbiology, 25(1):35–48, Jan. 2017.

14. Y. Fu, M. Pistolozzi, X. Yang, and Z. Lin. A comprehensive classification of coronaviruses and inferred cross-host transmissions. bioRxiv, 2020.

15. K. Y. Gorbunov and V. A. Liubetskii. Reconstructing genes evolution along a species tree. Molekuliarnaia Biologiia, 43(5):946–958, Oct. 2009.

16. D. E. Gordon, G. M. Jang, M. Bouhaddou, J. Xu, K. Obernier, K. M. White, M. J. O’Meara, V. V. Rezelj, J. Z. Guo, D. L. Swaney, et al. A sars-cov-2 protein interaction map reveals targets for drug repurposing. Nature, 583(7816):459–468, 2020.

17. R. L. Graham, J. S. Sparks, L. D. Eckerle, A. C. Sims, and M. R. Denison. Sars coronavirus replicase proteins in pathogenesis. Virus Research, 133(1):88–100, 2008. SARS-CoV Pathogenesis and Replication.

18. Y. Guan, B. Zheng, Y. He, X. Liu, Z. Zhuang, C. Cheung, S. Luo, P. Li, L. Zhang, Y. Guan, et al. Isolation and characterization of viruses related to the sars coronavirus from animals in southern china. Science, 302(5643):276–278, 2003.

19. B. Hie, E. D. Zhong, B. Berger, and B. Bryson. Learning the language of viral evolution and escape. Science, 371(6526):284–288, 2021.

20. E. Jacox, C. Chauve, G. J. Szollosi, Y. Ponty, and C. Scornavacca. eccetera: comprehensive gene tree-species tree reconciliation using parsimony. Bioinformatics, 32(13):2056, 2016.

21. I. Jungreis, R. Sealfon, and M. Kellis. Sars-cov-2 gene content and covid-19 mutation impact by comparing 44 sarbecovirus genomes. Nature Communications, 12(2642), 2021.

22. R. A. Khailany, M. Safdar, and M. Ozaslan. Genomic characterization of a novel sars-cov-2. Gene Reports, 19:100682, 2020.

23. D. Kim, J.-Y. Lee, J.-S. Yang, J. W. Kim, V. N. Kim, and H. Chang. The architecture of sars-cov-2 transcriptome. Cell, 181(4):914–921.e10, 2020.

24. M. Kordi and M. S. Bansal. Exact algorithms for duplication-transfer-loss reconciliation with non-binary gene trees. IEEE/ACM Transactions on Computational Biology and Bioinformatics, 16(4):1077–1090, 2019.

25. T. T.-Y. Lam, N. Jia, Y.-W. Zhang, M. H.-H. Shum, J.-F. Jiang, H.-C. Zhu, Y.-G. Tong, Y.-X. Shi, X.-B. Ni, Y.-S. Liao, W.-J. Li, B.-G. Jiang, W. Wei, T.-T. Yuan, K. Zheng, X.-M. Cui, J. Li, G.-Q. Pei, X. Qiang, W. Y.-M. Cheung, L.-F. Li, F.-F. Sun, S. Qin, J.-C. Huang, G. M. Leung, E. C. Holmes, Y.-L. Hu, Y. Guan, and W.-C. Cao. Identifying sars-cov-2-related coronaviruses in malayan pangolins. Nature, 583(7815):282–285, July 2020.

26. R. Libeskind-Hadas, Y.-C. Wu, M. S. Bansal, and M. Kellis. Pareto-optimal phylogenetic tree reconciliation. Bioinformatics, 30(12):i87–i95, 2014.

27. P. Liu, J.-Z. Jiang, X.-F. Wan, Y. Hua, L. Li, J. Zhou, X. Wang, F. Hou, J. Chen, J. Zou, and J. Chen. Are pangolins the intermediate host of the 2019 novel coronavirus (sars-cov-2)? PLOS Pathogens, 16(5):e1008421, May 2020.

28. K. S. Lole, R. C. Bollinger, R. S. Paranjape, D. Gadkari, S. S. Kulkarni, N. G. Novak, R. Ingersoll, H. W. Sheppard, and S. C. Ray. Full-length human immunodeficiency virus type 1 genomes from subtype c-infected seroconverters in india, with evidence of intersubtype recombination. Journal of Virology, 73:152–60, Jan 1999.

29. V. Makarenkov, B. Mazoure, G. Rabusseau, and P. Legendre. Horizontal gene transfer and recombination analysis of SARS-CoV-2 genes helps discover its close relatives and shed light on its origin. BMC Ecol Evo, 21(5), 2021.

30. D. P. Martin, B. Murrell, M. Golden, A. Khoosal, and B. Muhire. RDP4: Detection and analysis of recombination patterns in virus genomes. Virus Evolution, 1(1), 05 2015. vev003.

31. P. S. Masters, S. Perlman, et al. Coronaviridae, pages 825–858. Lippincott Williams & Wilkins Philadelphia, PA, 2013.

32. NCBI Resource Coordinators. Database resources of the National Center for Biotechnology Information. Nucleic Acids Research, 46(D1):D8–D13, Jan. 2018.

33. J. Á. Patiño-Galindo, I. Filip, M. AlQuraishi, and R. Rabadan. Recombination and lineage-specific mutations led to the emergence of sars-cov-2. bioRxiv, 2020.

34. M. Pérez-Losada, M. Arenas, J. C. Galán, F. Palero, and F. González-Candelas. Recombination in viruses: Mechanisms, methods of study, and evolutionary consequences. Infection, Genetics and Evolution, 30:296–307, Mar. 2015.

35. L. Pipes, H. Wang, J. P. Huelsenbeck, and R. Nielsen. Assessing Uncertainty in the Rooting of the SARS-CoV-2 Phylogeny. Molecular Biology and Evolution, 38(4):1537–1543, 12 2020.

36. D. F. Robinson and L. R. Foulds. Comparison of phylogenetic trees. Mathematical Biosciences, 53(1–2):131–147, 1981.

37. C. Scornavacca, E. Jacox, and G. J. Szöllosi. Joint amalgamation of most parsimonious reconciled gene trees. Bioinformatics, 31(6):841–848, 2015.

38. C. Scornavacca, W. Paprotny, V. Berry, and V. Ranwez. Representing a set of reconciliations in a compact way. Journal of Bioinformatics and Computational Biology, 11(02):1250025, 2013.

39. J. Sjostrand, A. Tofigh, V. Daubin, L. Arvestad, B. Sennblad, and J. Lagergren. A bayesian method for analyzing lateral gene transfer. Systematic Biology, 63(3):409–420, 2014.

40. A. Stamatakis. RAxML version 8: a tool for phylogenetic analysis and post-analysis of large phylogenies. Bioinformatics, 30(24451623):1312–1313, May 2014.

41. M. Stolzer, H. Lai, M. Xu, D. Sathaye, B. Vernot, and D. Durand. Inferring duplications, losses, transfers and incomplete lineage sorting with nonbinary species trees. Bioinformatics, 28(18):409–415, 2012.

42. M. A. Suchard, P. Lemey, G. Baele, D. L. Ayres, A. J. Drummond, and A. Rambaut. Bayesian phylogenetic and phylodynamic data integration using beast 1.10. Virus evolution, 4(29942656):vey016, 2018.

43. G. J. Szollosi, B. Boussau, S. S. Abby, E. Tannier, and V. Daubin. Phylogenetic modeling of lateral gene transfer reconstructs the pattern and relative timing of speciations. Proceedings of the National Academy of Sciences, 109(43):17513–17518, 2012.

44. G. J. Szollosi, E. Tannier, N. Lartillot, and V. Daubin. Lateral gene transfer from the dead. Systematic Biology, 62(3):386–397, 2013.

45. A. Tofigh. Using Trees to Capture Reticulate Evolution : Lateral Gene Transfers and Cancer Progression. PhD thesis, KTH Royal Institute of Technology, 2009.

46. A. Tofigh, M. T. Hallett, and J. Lagergren. Simultaneous identification of duplications and lateral gene transfers. IEEE/ACM Trans. Comput. Biology Bioinform., 8(2):517–535, 2011.

47. F. Tria, G. Landan, and T. Dagan. Phylogenetic rooting using minimal ancestor deviation. Nature Ecology and Evolution, 1:0193, 2017.

48. T. Wade, L. T. Rangel, S. Kundu, G. P. Fournier, and M. S. Bansal. Assessing the accuracy of phylogenetic rooting methods on prokaryotic gene families. PLOS ONE, 15(5):1–22, 05 2020.

49. F. Wu, S. Zhao, B. Yu, Y.-M. Chen, W. Wang, Z.-G. Song, Y. Hu, Z.-W. Tao, J.-H. Tian, Y.-Y. Pei, M.-L. Yuan, Y.-L. Zhang, F.-H. Dai, Y. Liu, Q.-M. Wang, J.-J. Zheng, L. Xu, E. C. Holmes, and Y.-Z. Zhang. A new coronavirus associated with human respiratory disease in china. Nature, 579(7798):265–269, Mar. 2020.

50. T. Zhang, Q. Wu, and Z. Zhang. Probable pangolin origin of sars-cov-2 associated with the covid-19 outbreak. Current Biology, 30(7):1346–1351, 2020.

51. P. Zhou, X.-L. Yang, X.-G. Wang, B. Hu, L. Zhang, W. Zhang, H.-R. Si, Y. Zhu, B. Li, C.-L. Huang, H.-D. Chen, J. Chen, Y. Luo, H. Guo, R.-D. Jiang, M.-Q. Liu, Y. Chen, X.-R. Shen, X. Wang, X.-S. Zheng, K. Zhao, Q.-J. Chen, F. Deng, L.-L. Liu, B. Yan, F.-X. Zhan, Y.-Y. Wang, G.-F. Xiao, and Z.-L. Shi. A pneumonia outbreak associated with a new coronavirus of probable bat origin. Nature, 579(7798):270–273, Mar. 2020.

